# Foliar salt spray exclusion and tissue tolerance underlie local adaptation to oceanic salt spray

**DOI:** 10.1101/2025.10.16.682946

**Authors:** Madison L. Plunkert, P. C. Durant, Katherine Egeler, David B. Lowry

**Affiliations:** Department of Plant Biology, 612 Wilson Road, Michigan State University, East Lansing, MI, 48824, USA; Ecology, Evolution, and Behavior Program, 567 Wilson Road, Michigan State University, East Lansing, MI, 48824, USA

**Keywords:** abiotic stress, adaptation, coastal, *Erythranthe guttata*, leaf surface traits, *Mimulus guttatus*, salt spray, yellow monkeyflower

## Abstract

**Premise:** Surviving under oceanic salt spray is critical for plants in coastal ecosystems, yet the mechanisms of coastal plant resilience to salt spray are not well understood. We investigated mechanisms of salt spray adaptation by comparing five latitudinal pairs of yellow monkeyflower accessions locally adapted to coastal and inland habitats.

**Methods:** We measured sodium levels in coastal and inland leaves exposed to experimental salt spray in the lab, and compared leaf surface traits that may contribute to differences in sodium uptake between ecotypes. We assayed tissue tolerance by allowing sodium to enter leaves through wounding and recording time to necrosis for coastal and inland accessions.

**Key Results:** Coastal monkeyflowers take up less sodium through the leaf surface under experimental salt spray, which may contribute to local adaptation under oceanic salt spray in coastal habitats. Higher water content per unit leaf area and less water loss under salt spray further reduce sodium concentrations in salt-sprayed coastal leaves compared to inland counterparts. The coastal ecotype also shows greater tissue tolerance to sodium than the inland ecotype.

**Conclusions:** Coastal monkeyflowers employ salt spray exclusion and tissue tolerance mechanisms for salt spray resilience. Our results suggest that plant adaptation to coastal habitats may often involve the evolution of multiple mechanisms to survive stress imposed by oceanic salt spray.

## INTRODUCTION

Along the vast coastlines of the Earth, oceanic salt spray plays a key role in structuring the species composition of plant communities (Boyce, 1954; Barbour, 1978; Du and Hesp, 2020). These coastal communities experience oceanic salt spray not only as an abiotic stressor affecting species distribution, with many plant species unable to survive the intense salt spray closest to the coast, but also as an agent of selection that drives local adaptation (Popovic and Lowry, 2020). Despite the influence of salt spray resilience on which plant species occur in coastal habitats, we know comparatively little about how populations *within* a species locally adapt to salt spray (Du and Hesp, 2020). Uncovering the traits that influence resilience to abiotic stressors like salt spray is a critical goal of plant adaptation research. By comparing salt spray-sensitive and salt spray-adapted populations of a single species, we can determine the traits, physiological processes, and genetic mechanisms that underlie local adaptation to salt spray.

Salt spray has long been suspected to play a key role in adaptive, genetic, morphological, and physiological differences between coastal and nearby inland populations. Many distantly related plant species contain both upright, early flowering, inland forms and prostrate, late-flowering, coastal forms (Turesson, 1922; Clausen and Hiesey, 1958). Over seventy years ago, Boyce hypothesized that the shorter stature of the coastal populations within species was the result of adaptations to oceanic salt spray and suggested that these coastal forms each be referred to as a “salt spray ecotype” (Boyce, 1954). More recently, research on *Senecio lautus* Willd. found that coastal headland populations display prostrate growth and agravitropism, whereas those from the more sheltered dune sites grow upright (Wilkinson et al., 2021).

Similarly, compact coastal *Setaria viridis* L. are more resilient to salt spray than their tall inland counterparts (Itoh, 2021; Itoh et al., 2024). Phenology also contributes to salt spray escape; in the yellow monkeyflower *Mimulus guttatus* DC. (Phrymaceae; syn. *Erythranthe guttata* DC., Barker et al., 2012; Lowry et al., 2019b), coastal plants flower late in the season, coinciding with calmer wind speeds, while inland plants flower during high spring winds, leading to intense salt spray exposure in transplant experiments (Hall and Willis, 2006; Zambiasi and Lowry, 2024; Toll et al., 2025). While the aforementioned studies suggest a widespread salt spray escape mechanism, the strategies coastal plants use to cope with salt spray after it contacts the plant remain unknown.

In addition to the scarcity of evolutionary studies on salt spray, we also lack an understanding of salt spray response as a physiological and cellular process. When salt reaches the roots of a salt-tolerant plant via saline soils, that plant may survive through osmotic tolerance, sodium exclusion, tissue tolerance mechanisms, or a combination of the three (Munns and Tester, 2008; Rajendran et al., 2009). Other studies categorize salinity responses as escape, avoidance, tolerance, and recovery (Grewell et al., 2025). Comparable frameworks do not exist for salt spray. We suggest that three non-mutually exclusive mechanisms may underlie salt spray resilience: **salt spray escape, salt spray exclusion,** and **tissue tolerance** (Box 1).

We propose that salt spray escape mechanisms employ features of plant architecture to prevent oceanic salt spray from contacting the shoot (Figure 1A). Salt spray escape is the best supported mechanism of salt spray adaptation, but other mechanisms also exist. Under experimental salt spray, where *Setaria viridis* plants were exposed to salt spray regardless of height, coastal plants sustained less injury than inland plants (Itoh et al., 2024), suggesting other adaptive mechanisms in addition to salt spray escape that occur once salt spray has contacted the plant.

**Figure 1.**
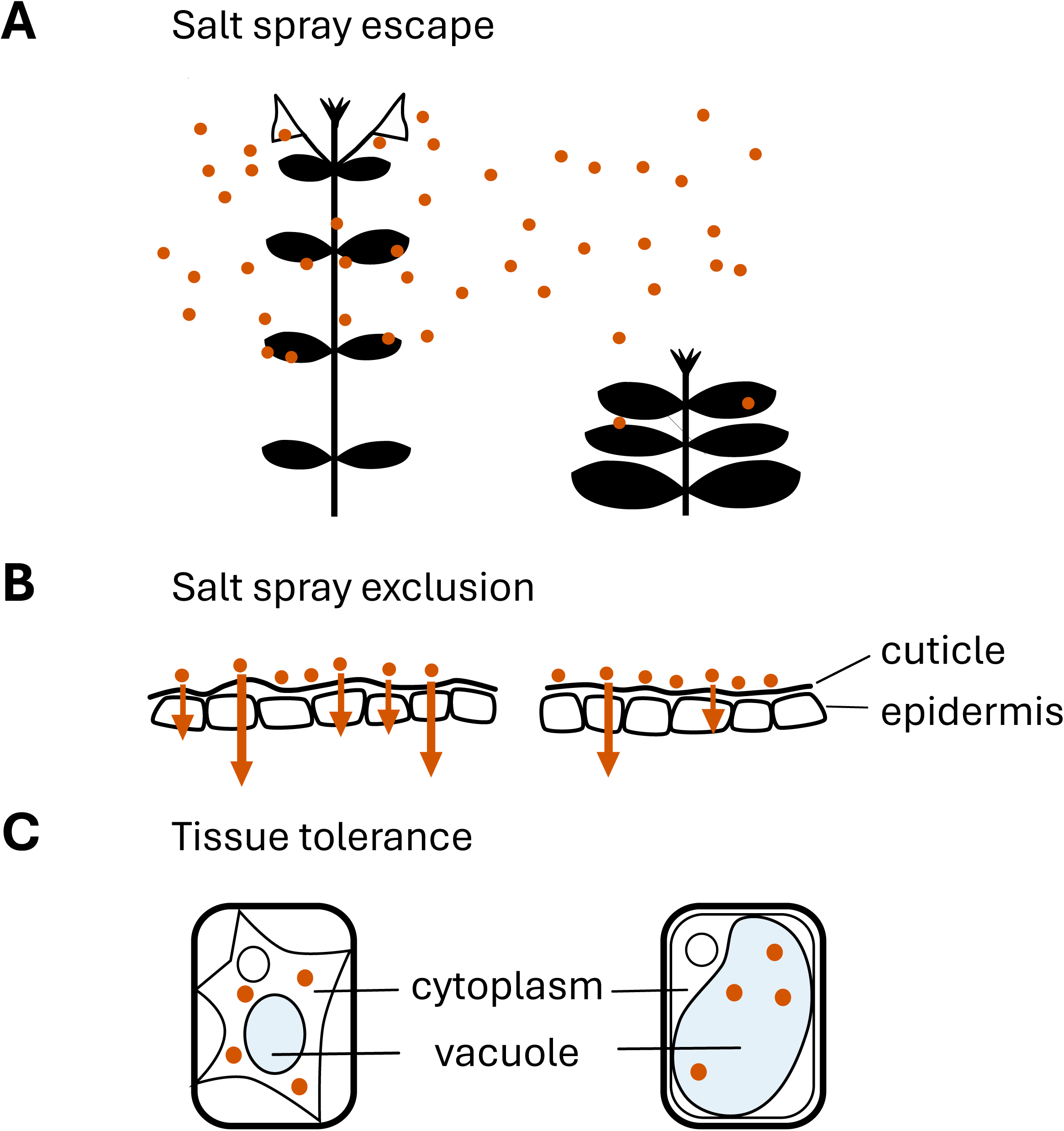
Conceptual model showing mechanisms of adaptation to salt spray, with a salt spray-sensitive inland plant on the left and a salt spray-adapted coastal plant on the right. (A) Salt spray intensifies with height, such that tall inland plants are more exposed to salt spray than short coastal plants. Inland ecotype flowering time corresponds with time periods of high spray intensity, while coastal ecotype flowering time is delayed until spray intensity is lower. (B) Salt on the leaf surface enters the leaf tissue at a greater rate in inland than coastal plants. (C) Inland leaves have lower tissue tolerance to salt inside cells than coastal leaves. Coastal leaves have high succulence and they better maintain this succulence under salt spray, which may help them sequester sodium in large vacuoles.

If salt spray contacts the shoot, plant traits may prevent salt from entering the tissue via salt spray exclusion (Figure 1B). In this mechanism, characteristics of the leaf surface such as leaf wettability or stomatal traits reduce entry of salt spray. Wettable (or hydrophilic) leaves tend to take up less water from the leaf surface than non-wettable leaves (Ahmad and Wainwright, 1976; Cavallaro et al., 2022). Stomata present another opportunity for seawater to enter leaf tissue (Burkhardt et al., 2012; Emery, 2016; Berry et al., 2019; Guzmán-Delgado et al., 2021). Many studies show that plants with fewer open stomata have less foliar water or sodium uptake (Grieve and Pitman, 1978; Burkhardt et al., 2012; Guzmán-Delgado et al., 2021). Further, amphistomy, the placement of stomata on both sides of a leaf, can offer a photosynthetic advantage to placing stomata only on the abaxial surface (Triplett et al., 2024). However, amphistomy may have unexplored consequences under salt spray if seawater enters through exposed adaxial stomata. Thus, stomatal anatomy and other salt spray exclusion traits potentially contribute to plant resilience under salt spray.

The latest-acting proposed mechanism for survival under salt spray is tissue tolerance, in which salt spray contacts the leaf and enters the tissue, but the plant maintains homeostasis and prevents cellular damage (Figure 1C). Sodium accumulation in leaves contributes to plant mortality under both soil salinity and salt spray, allowing us to draw on the broader salt tolerance literature (Grieve and Pitman, 1978; Haq et al., 2013). To cope with excess sodium, leaf cells may sequester sodium in the vacuole, preventing it from disrupting cytosolic enzyme function (Apse et al., 2003; Munns and Tester, 2008). Finally, succulent tissues can prevent sodium from reaching lethal concentrations simply by diluting it through their high water content (Liu et al., 2024). Plants leverage diverse mechanisms to escape, exclude, or tolerate salt spray stress.

To understand adaptive mechanisms of survival under salt spray, we leverage genetic variation in the yellow monkeyflower *M. guttatus,* a model system for the genetic basis of local adaptation. *M. guttatus* has extensive genetic diversity, evidence of local adaptation, and genomic resources, increasing our understanding of the phenotypic and genetic variation underlying local adaptation (Twyford and Friedman, 2015; Lovell et al., 2025). The inland annual and coastal perennial ecotypes are locally adapted to their respective habitats (Lowry et al., 2008; Lowry and Willis, 2010). The inland ecotype is upright in growth form, earlier flowering, and generally taller than the coastal ecotype, which contributes to the inland ecotype’s high mortality under salt spray in reciprocal transplant experiments (Toll et al., 2025, Appendix S1). When inland annuals are transplanted to the coastal environment, they can survive only when protected from salt spray and aboveground herbivores (Popovic and Lowry, 2020). In unprotected plants, salt spray is the major cause of mortality for inland annuals at coastal sites (Toll et al., 2024), while coastal perennials at inland sites are killed by a terminal summer drought (Hall and Willis, 2006; Lowry and Willis, 2010). Intriguingly, this shift in inland annual mortality rates at coastal sites (from 51% outside to 3% inside the exclosure) occurs without manipulating belowground factors (Popovic and Lowry, 2020). This result indicates that aboveground salt spray, not soil salinity, is a key factor driving local adaptation in coastal habitats. Comparing plant traits and responses to salt spray between the salt spray-sensitive inland annual and the salt spray-adapted coastal perennial allows us to identify mechanisms for adaptation to salt spray.

Salt spray escape, through plant stature and phenology, is an important strategy in coastal monkeyflower, but this strategy alone does not fully account for increased survival of the coastal ecotype under salt spray. Coastal plants outperformed inland plants even when tall and flowering, indicating that height and flowering time are not the only mechanisms of salt spray adaptation (Toll et al., 2025). This observation suggests that later-acting mechanisms after salt spray contacts plant tissues are also important for adaptation in this system.

In this study, we test whether multiple resilience strategies underlie salt spray adaptation in coastal yellow monkeyflowers. To compare the effectiveness of all salt exclusion barriers in the coastal and inland ecotypes, we measured sodium levels in leaves treated with salt spray or deionized water in the lab. We tested potential salt exclusion mechanisms by measuring leaf surface traits in coastal and inland leaves that may affect saltwater entry into leaf tissue. Finally, we compared tissue tolerance in coastal and inland monkeyflowers by measuring succulence, a predicted tissue tolerance trait, and by exposing leaf discs to salt and recording the time to tissue death. Broadly, the study investigates the role of salt spray exclusion and tissue tolerance mechanisms in salt spray resilience in coastal monkeyflower, addressing the role of multiple strategies in surviving a major selective agent of coastal ecosystems.

## MATERIALS AND METHODS

### Plant material and growth conditions

We performed all experiments using inland annual and coastal perennial accessions derived from the species complex of the yellow monkeyflower *Mimulus guttatus,* which is found across western North America, often in seep habitat. They are bee-pollinated and primarily outcrossing, and many perennials are capable of clonal propagation through stolons or rhizomes (Willis, 1993; Coughlan et al., 2021). Some taxonomic treatments of these ecotypes regard them as separate species with the coastal perennial ecotype being separated out as the magnificent monkeyflower *M. grandis* (Greene) and the inland annual ecotype being separated out as *M. microphyllus* (Benth.) (Barker et al., 2012; Nesom, 2014). While the distinct population structure of the magnificent monkeyflower is supported by population genetic analyses, the separation of *M. microphyllus* as a group from the broader species complex is contradicted by genetic data (Oneal et al., 2014; Twyford and Friedman, 2015; Frayer et al., 2026). These coastal perennials and inland annuals are genetically diverged, yet readily form viable hybrids, harbor more genetic variation between populations within ecotypes than between ecotypes (Lowry et al., 2008), and have few fixed genetic differences (Gould et al., 2017). Given their population structure and incomplete reproductive isolation, we consider them as ecotypes of *M. guttatus* for the purpose of this study.

To establish whether coastal and nearby inland populations differ across the geographic range of the coastal ecotype (From Point Conception in California to the Columbia River region of Oregon and Washington states), we selected five coastal perennial accessions and five nearby inland annual accessions from similar latitudes of origin for this study (Figure 2, Appendix S2). Accessions had been previously inbred in the lab for 1–7 generations based on the availability of seed stocks, and we selected early-generation inbred lines when possible. Using these lines allowed us to avoid potential bias in the results due to maternal effects of seed maturation occurring in the coastal and inland environments. Plants were grown in a growth chamber (model GRC-40, BioChambers Inc., Winnipeg, Manitoba, Canada) with a 12-hour photoperiod, 22°C day and 18°C night temperatures, 60% relative humidity, and a light intensity of 460 µmol m^-2^ s^-1^ photosynthetic active radiation. Plants were watered by subirrigation every other day. For ion measurements and stomatal conductance measurements, a separate batch of plants were grown for 3 months to a large rosette stage in a growth chamber (model TPC-37, BioChambers Inc., Winnipeg, Manitoba, Canada) and large, undamaged leaves were selected for spray treatment and subsequent ion measurements.

**Figure 2.**
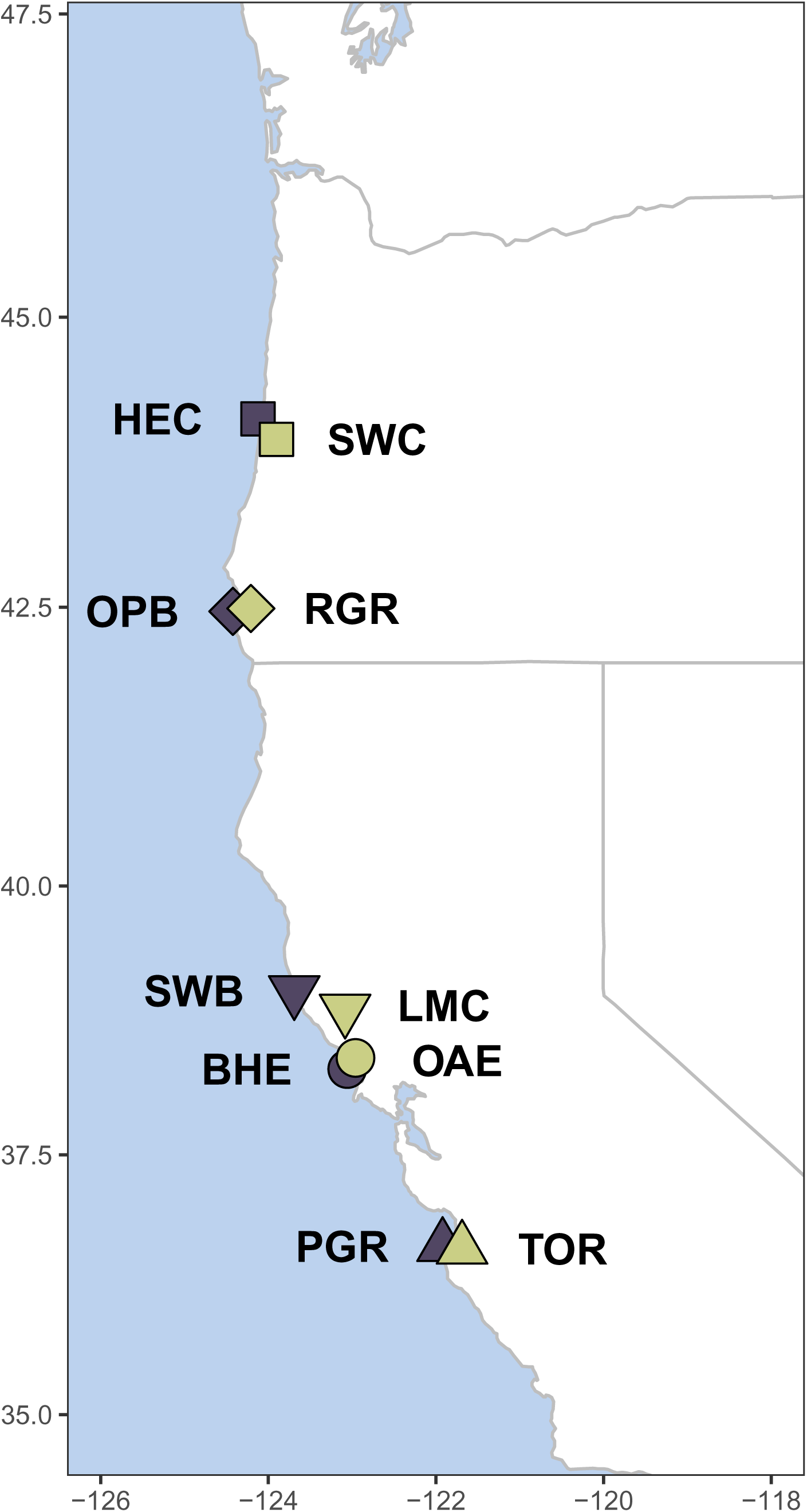
Map of sites of origin for accessions used in this study, with coastal accessions in gray-purple, inland accessions in yellow, and shapes indicating which coastal and inland accessions are latitudinally paired for later analysis. Shapes and colors used for each accession are consistent throughout Appendices, where accession-specific data are shown. More accession details can be found in Appendix S2.

### Salt exclusion test

To test whether coastal and inland monkeyflowers accumulate different levels of sodium when exposed to foliar NaCl, we sprayed plants from all accessions with either 0.63 M NaCl (ocean concentration) solution or deionized water using an airbrush (model G222, Master Airbrush, North Las Vegas, Nevada, USA) with a 0.3 mm^2^ needle at 207 kPa pressure, holding the airbrush about 15 cm away from the adaxial surface of the targeted leaf and spraying for five seconds. We performed two spray treatments 18 hours apart, allowed leaves to dry for two hours after the second treatment, and then sampled four water-treated and four salt-treated leaves from each of the 10 accessions. This short time interval between salt spray treatment and leaf collection allowed us to make comparisons between relatively healthy leaf tissue while using sodium chloride solutions at a realistic ocean concentration, since inland annuals show tissue necrosis in the first few days of salt spray (Lowry et al., 2008, 2009). We observed some turgor loss but no necrosis which can substantially increase sodium uptake (Munns et al., 2016). The four leaves were sampled from 2–3 different plants and we treated leaves as biological replicates. We immediately recorded fresh mass of the collected leaves, scanned them with a document scanner for leaf area determination with ImageJ (version 1.54f, Schneider et al., 2012), rinsed them in 0.5 M EDTA (pH = 8), and rinsed them twice in ultrapure water (produced by Milli-Q® system, Millipore Sigma, St. Louis, Missouri, USA). We dried the leaves, ground them into a powder, and degassed and mineralized the powder by combining 1 mg of powdered leaf with 1 mL of 3 parts 65% nitric acid to 1 part hydrogen peroxide and incubating for 24 hours at room temperature, then overnight at 85°C. We combined the mineralized solution with 4 mL of ultrapure water, and used the resulting solution for microwave plasma atomic emission spectroscopy (MP-AES) with the 4210 Agilent MP-AES instrument (Agilent Technologies, Inc., Santa Clara, California, USA). Sodium ion concentrations were determined from a standard curve (Appendix S3) with Agilent MP Expert software (Agilent Technologies, Inc., Santa Clara, California, USA). We used the mean ion concentration in ppm of three technical replicates on each sample for downstream analysis. We verified that the rinsing step effectively removed sodium ions on the leaf surface, leaving behind the sodium inside the leaf tissue, by comparing sodium concentrations of leaves that were salt-sprayed and rinsed immediately to those that had salt spray sit on the leaf surface for 18 hours (Appendix S4). Based on this test of rinsing residual surface salts, we proceeded with MP-AES analysis of rinsed leaves as an approach to measuring sodium concentrations inside salt-sprayed leaf tissue.

To assess salt spray exclusion, we calculated µmol Na^+^ per unit leaf area and molarity of Na^+^ in the leaf tissue by dimensional analysis using the fresh mass, dry mass, and area of each leaf, and the density of mineralization solution (1.062 g/mL). We calculated succulence as (fresh weight – dry weight)/leaf area for the same leaves, following Guo et al., 2023, and leaf mass per area (LMA) as leaf dry mass (including leaf petioles) divided by leaf area.

### Investigation of potential leaf surface traits for salt spray exclusion

To test whether leaf traits may contribute to sodium exclusion, we measured leaf surface traits in leaves from 15 individuals of each of the five coastal and five inland accessions. We arranged the plants haphazardly in a growth chamber and phenotyped them when the third pair of true leaves had begun to expand. Measurements were taken on well-watered plants before the lights came on in the chamber.

#### Leaf wettability phenotyping

To test for differences in leaf wettability between ecotypes, we measured the contact angle of a droplet of water on the surface of the leaf. Leaves with low contact angle values are more wettable (or less hydrophobic) than leaves with high contact angle values (Ahmad and Wainwright, 1976; Cavallaro et al., 2022). We cut a leaf from the first pair of true leaves in half along the midvein to determine the contact angle of the adaxial and abaxial surfaces, sampling one leaf from each of 15 plants from each accession. We affixed each half to a flat surface using double-sided tape and placed a 10 µl droplet of distilled water on the surface under study. Pilot studies showed that contact angles are similar between distilled water and NaCl solution. We photographed the droplet from the side using a Nikon D3400 digital camera with AF-S Micro NIKKOR 85 mm 1:3.5G ED lens (Nikon, Tokyo, Japan) and repeated for the other leaf surface. We measured contact angle using the Contact Angle plugin (Brugnara, 2006) in ImageJ (version 1.54i, Schneider et al., 2012) using an elliptical fit.

As a second assay of leaf wettability, we performed a leaf water drop adhesion assay modified from (Wang et al., 2014). We cut an undamaged leaf from the first or second true leaf pair from each of 15 plants from each accession and recorded fresh leaf mass. After submerging each leaf in water for 10 seconds, we held the leaf suspended in the air for 10 seconds to allow water to run off, then weighed the leaf again to assess how much water had clung to the surface. To determine leaf area, we scanned leaves along with a ruler on a document scanner at 300 dpi and measured leaf area in ImageJ (version 1.54f, Schneider et al., 2012) by setting the scale according to the ruler, converting the image to 8-bit format, manually adjusting the threshold to correspond to the entire leaf, and selecting the leaf area. We calculated water retained after submergence per unit leaf area as (mass of submerged leaf mass – fresh mass)/(leaf area*2), multiplying leaf area by 2 since water can interact with both sides of the leaf.

Leaves that retained more surface water per unit leaf area are more wettable than those that retained less water per unit leaf area.

#### Stomatal size and density phenotyping

To compare stomatal traits between coastal and inland accessions, we cut a leaf from the second pair of true leaves from each of 15 plants from each accession in half along the midvein and applied clear nail polish to the adaxial surface of one half and the abaxial surface of the other half to make stomatal peels. We imaged stomatal peels using an AmScope compound microscope with 100x magnification (AmScope, Irvine, California, USA), Nikon microscope camera, and Nikon BR software (Nikon, Tokyo, Japan) with a field of view of 0.9437 mm^2^. We manually counted stomata in 1–3 fields of view per stomatal peel. We measured stomatal length as the straight-line distance from one end of the guard cells to the other on 10 stomata in one field of view for both sides of one leaf per plant using ImageJ (version 1.54g, Schneider et al., 2012). We used the mean of the 10 measured stomata on each leaf for downstream analysis. We used stomatal length and stomatal density to estimate the fraction of leaf epidermis area that was allocated to stomata following Muir et al. (2023). Finally, we calculated amphistomy, the production of stomata on both surfaces of a leaf, as adaxial stomatal stomatal count / (adaxial stomatal count + abaxial stomatal count) in the same size field of view, to test for differences in the placement of stomata on the salt spray-exposed adaxial surface and protected abaxial surface between ecotypes.

#### Stomatal conductance measurements

To determine whether coastal and inland leaves differ in the opportunity for saltwater uptake via stomata, we measured stomatal conductance in the coastal and inland accessions. We measured stomatal conductance to water (g*_sw_*, in mol m^-2^s^-1^) using a LI-600 porometer (LI-COR, Lincoln, Nebraska, USA) on the adaxial surface of 6 leaves sampled from 2–3 mature plants for each accession, treating leaves as biological replicates. Measurements were completed between 1–3 hours after lights came on in the chamber, with genotypes measured in a haphazard order. We measured stomatal conductance on a different set of plants from the stomatal density, stomatal size, and wettability traits.

#### Succulence and LMA

To characterize ecotypic differences in succulence and LMA with higher replication within genotypes, we calculated succulence and LMA using the same calculation method as the salt exclusion test. The same leaves used in the leaf water drop adhesion assay (one leaf from each of 15 plants per accession) were dried to get leaf dry mass for succulence and LMA.

### Test of tissue tolerance

To test whether coastal accessions prevent damage better than inland accessions once salt enters the leaf, we compared time to tissue death in leaf discs exposed to salt. In this assay, sodium chloride can enter the leaf through the wound at the edge of the leaf disc, so tissue tolerance mechanisms, but not leaf surface properties that prevent salt entry, can influence time to death. We prepared plates of half-strength Murashige and Skoog Basal Medium with Gamborg′s vitamins (M0404-10L, Sigma-Aldrich, Inc., St. Louis, Missouri, USA) and agar (PTP01-1KG, Caisson Labs, Smithfield, Utah, USA), adjusted to pH 5.6 with KOH, and either supplemented with 100 mM NaCl (salt) or not (control) for a total of 10 plates. Two leaf discs from each accession were arranged randomly onto each plate with the abaxial side on the media, since necrosis is easier to visually distinguish from the green adaxial surface than the red abaxial surface seen in some genotypes. The plates were then stored at 20°C under constant light. Each leaf disc was observed and photographed daily until all discs were fully necrotic. The day when necrosis began, as well as the day the disc fully necrotized, was recorded for each leaf disc by an observer who did not know which disks were coastal and inland.

We excluded disks where algal growth prevented us from scoring necrosis, leaving 6–10 disks per accession per treatment for the final analysis.

### Statistical analysis

For experiments involving salt and control treatments, we fit a general linear mixed model to model the fixed effects of ecotype, treatment, and their interaction, and the random effect of accession nested within latitudinal pair, on the response variable of interest (response variable ∼ ecotype*treatment + (1| latitudinal pair / accession). For one response variable, concentration of sodium ions, including latitudinal pair in the random effects led to a model convergence problem (non-positive-definite Hessian matrix). We dropped latitudinal pair and included only accession in the random effects, which led to negligible change in effect sizes and significance.

For all leaf surface traits examined (contact angle, stomatal density, stomatal size, stomatal conductance, estimated fraction of area allocated to stomata, leaf water drop adhesion assay), we fit a general linear mixed model to model the fixed effect of ecotype and the random effect of accession nested within latitudinal pair on the leaf trait of interest (leaf trait ∼ ecotype + (1 | latitudinal pair / accession + ecotype). For one response variable, abaxial stomatal density, including latitudinal pair in the random effects led to a model convergence problem (non-positive-definite Hessian matrix). We dropped latitudinal pair and included only accession in the random effects for that model, which led to negligible change in effect sizes and significance.

General linear mixed models were fit using the *glmmTMB* package, version 1.1.11 (Brooks et al., 2017) and visualized as least-squares means (LSMs) for each ecotype, or each combination of ecotype and treatment, from the *emmeans* package (version 1.11.0, Lenth et al., 2025). To visualize differences between latitudinally paired accessions, we plotted accession arithmetic means and standard errors of the mean. All statistical analyses were conducted in R version 4.4.1 (R Core Team, 2024).

## RESULTS

### Salt spray exclusion

Following the saltwater spray application, inland plant leaves had a 2.8 times higher internal concentration of sodium than coastal plant leaves (treatment*ecotype interaction = 0.29 M, *P* < 0.0001, Figure 3A, Appendix S5). Because coastal and inland plants had such different sized leaves, it was critical that we calculated sodium entry on a per unit leaf area basis to test whether the masses of internal leaf sodium reflected true differences in salt spray exclusion. Indeed, our per leaf area calculation demonstrated that coastal plant leaves exclude sodium better than inland leaves (treatment*ecotype interaction = 2.66 mol/cm^2^, *P* = 0.0014, Figure 3B, Appendix S5).

**Figure 3.**
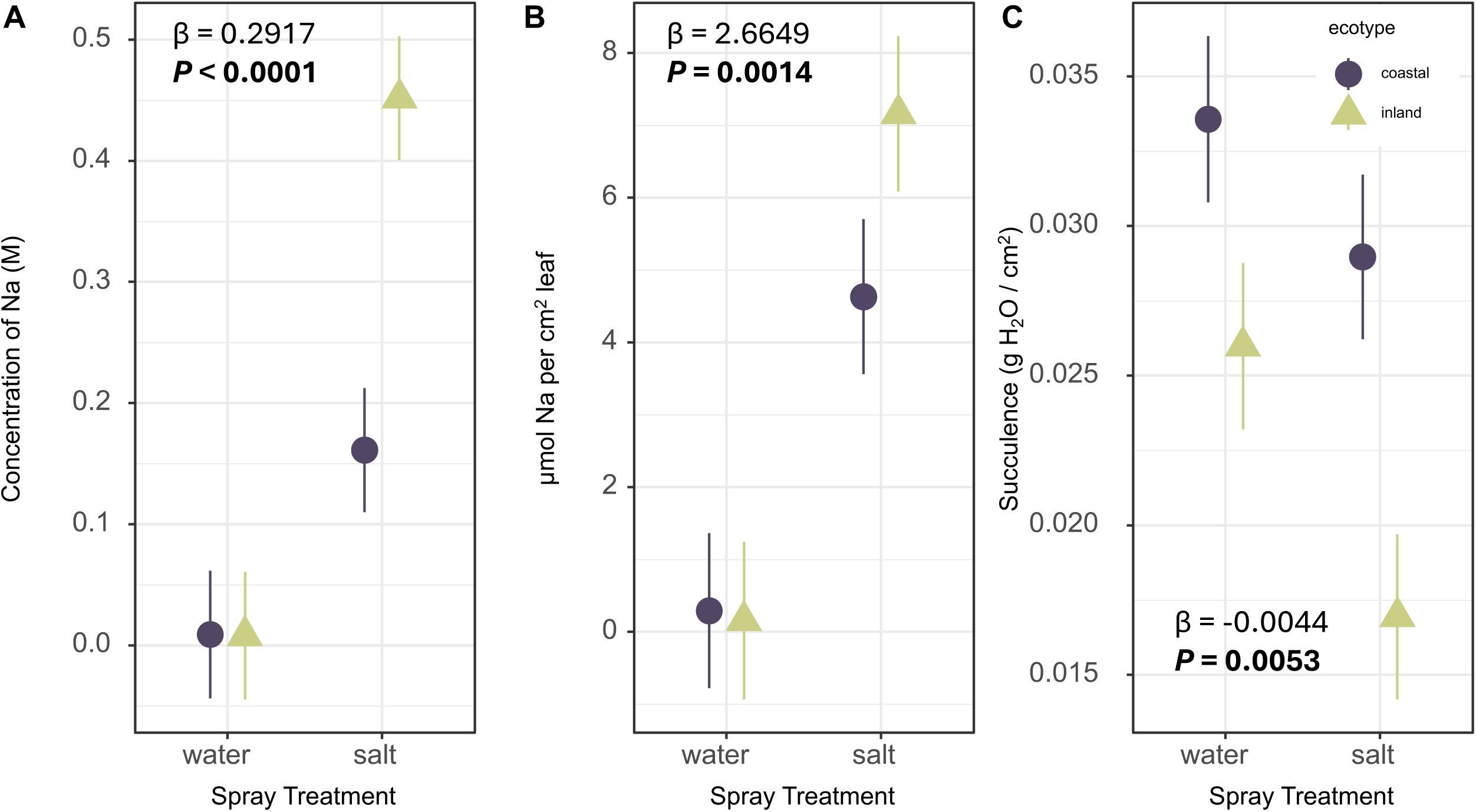
Sodium and water levels in coastal and inland ecotypes after salt spray or control treatment, with LSMs from general linear mixed models for each ecotype under salt or water treatment. Error bars show 95% confidence intervals. β and p-values indicate the interaction effect of inland ecotype and salt treatment. (A) Sodium concentration (M) for each ecotype and treatment after spray, assuming all Na is in solution in the volume of water in the leaf at the time of sampling. (B) µmol of Na per unit leaf area for each ecotype and treatment. (C) Succulence as (fresh mass - dry mass) / leaf area, in grams H_2_O per cm^2^ leaf for each ecotype and treatment.

The finding of salt spray exclusion on a per-area basis was broadly consistent among accessions, except for the coastal accession HEC which had inland-like levels of sodium per unit area (Appendix S6). Interestingly, coastal populations vary in whether they reached this reduced sodium concentration by reduced sodium, greater water content, or both (Appendix S6). Further, coastal leaves were more succulent (as mass of water per unit leaf area) and lost less water when salt sprayed than inland leaves (treatment*ecotype interaction β = -0.0044 g/cm^2^, *P* = 0.0053, Figure 3C, Appendix S5). Since both sodium and water content influenced the molarity of sodium, the sodium entry per unit leaf area and the water per unit leaf area together drove the large difference in sodium concentration in salt-sprayed coastal and inland leaves.

### Salt spray effects on leaf mass per area

In addition to the changes in sodium concentration mediated by sodium exclusion and water content, the leaf mass per area (LMA) of coastal perennials and inland annuals also responded differentially to salt spray treatment. Both ecotypes had greater LMA in salt-sprayed than water-sprayed leaves, with inland annuals experiencing a greater increase in LMA when salt-treated (ecotype*treatment interaction effect = 14.811 g m^-2^, *P* = 0.0002, Figure 4, Appendices S7, S8). Notably, the increase in LMA (19.5 g m^-2^ higher in salt-sprayed inland than water-sprayed inland leaves) was nearly five-fold larger than the mass of sodium chloride (7 μmol/cm^2^) in salt-sprayed inland leaves (Figure 3B; calculation assuming equimolar sodium and chloride uptake). We detected significant effects of ecotype, but not treatment nor the interaction of ecotype and treatment, on leaf area and dry mass alone. In summary, both ecotypes increased LMA with salt spray, with a more dramatic increase in inland annuals, and the source of this increase was unknown.

**Figure 4.**
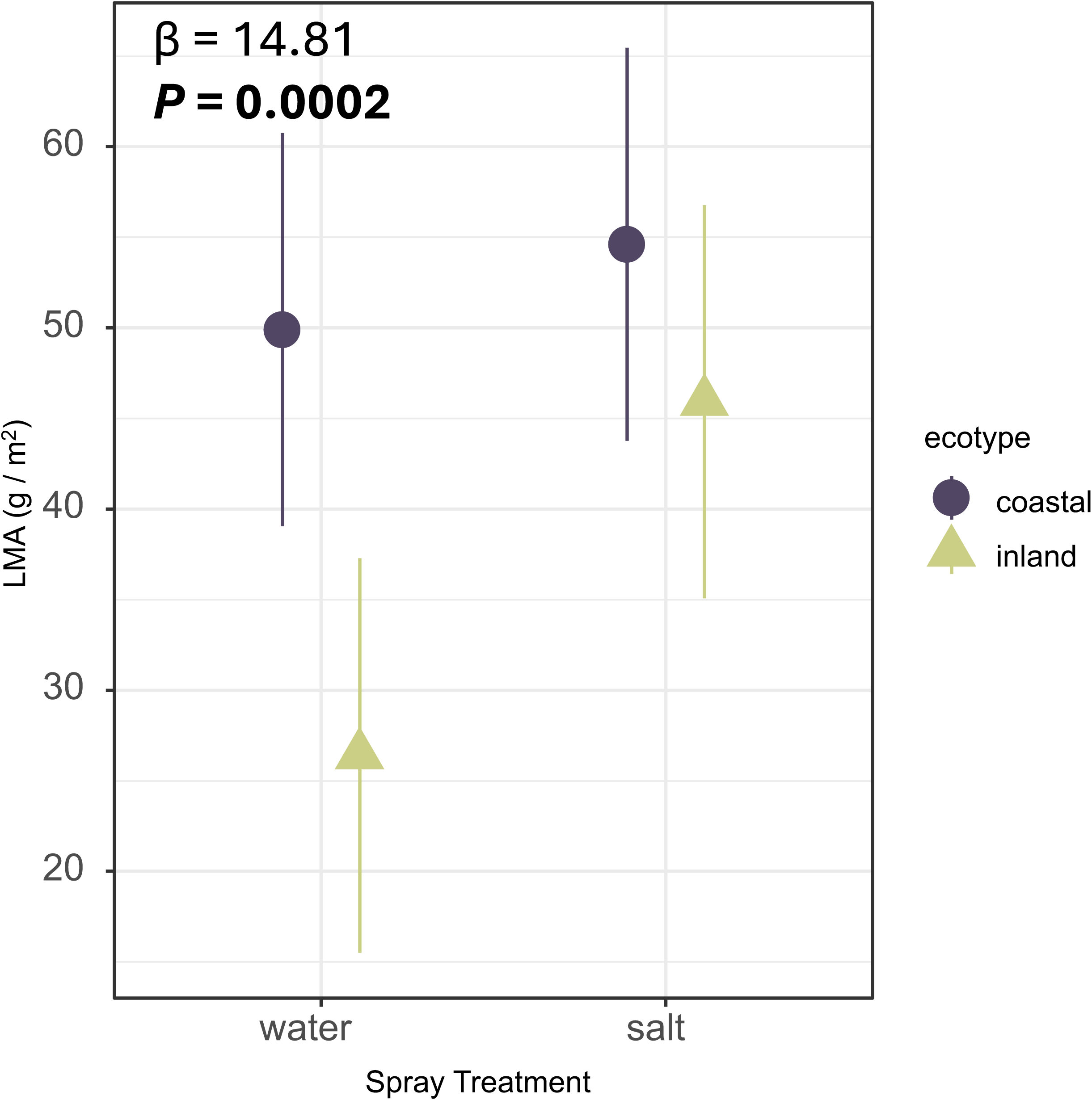
Dry leaf mass per area as g/m^2^ (y-axis) for each ecotype under water or salt spray treatment (x-axis). Each point indicates LSMs from general linear mixed models for each ecotype under salt or water treatment, with four leaves measured per accession for each treatment. Error bars show 95% confidence intervals. β and *P*-value indicate the interaction effect of inland ecotype and salt treatment.

### Leaf wettability

If leaf wettability were an adaptive mechanism underlying greater salt spray exclusion in the coastal ecotype, we would expect coastal leaves to be less wettable. Prior studies have shown that less wettable leaves cause saltwater to bead up and roll off, instead of staying on the leaf surface where it could enter the tissues through the stomata or cuticle (Ahmad and Wainwright, 1976; Humphreys et al., 1986; Lenz et al., 2022).

However, we found that there was not a significant difference in contact angles of water on the leaves of the inland and coastal ecotypes for both the adaxial (inland ecotype effect = 4.48°, *P* = 0.19) or abaxial (inland ecotype effect = 5.02°, *P* = 0.060) surfaces Figure 5A, 5B, Appendices S9, S10). We did find that the leaves of the inland ecotype retained less water on the surface per unit leaf area than the coastal ecotype in the leaf water drop adhesion assay (inland ecotype effect = -0.0061 g H_2_O cm^-2^, *P* = 0.0003, Figure 5C, Appendices S9, S10). Together, these assays indicated that the leaves of the inland and coastal ecotypes have similar levels of wettability, with inland leaves having less wettable leaves than coastal plants for some metrics. Reduced leaf wettability is therefore unlikely to contribute to salt spray adaptation by salt spray exclusion in coastal monkeyflower.

**Figure 5.**
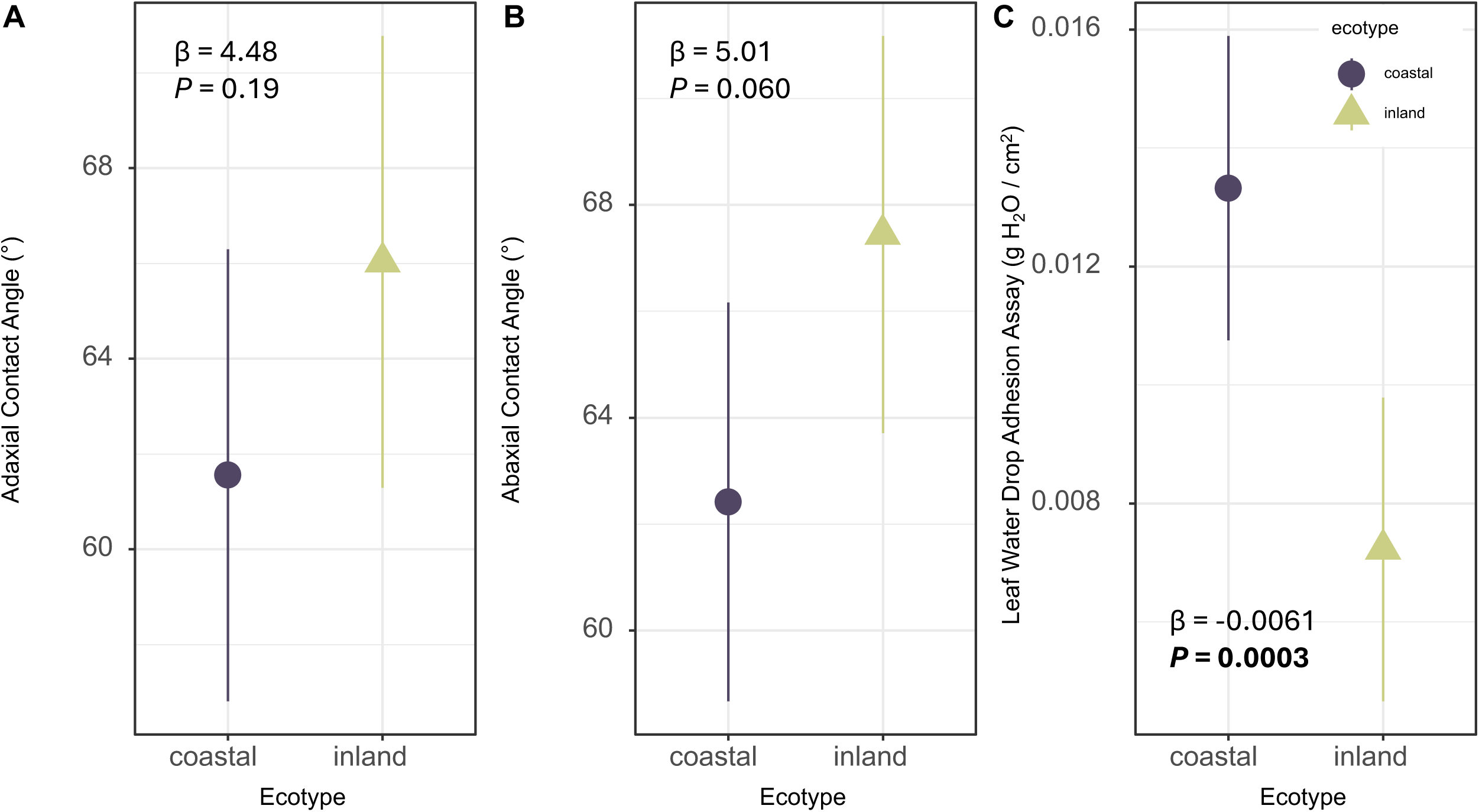
Comparison of leaf wettability between ecotypes. Points indicate LSMs for each ecotype from general linear mixed models, with 15 individuals measured per accession, and error bars show 95% confidence intervals. β and p-value indicate the effect of inland ecotype on the response variable. (A) Contact angle of a drop of water on the adaxial surface in degrees (y-axis) against ecotype (x-axis). (B) Contact angle of a drop of water on the abaxial surface in degrees (y-axis) against ecotype (x-axis). (C) Leaf water drop adhesion assay result in grams H_2_O on the surface of a submerged leaf per cm^2^ of leaf area (y-axis) against ecotype (x-axis).

### Stomatal traits

Stomatal density on both the adaxial (inland ecotype effect = -87.82 mm^-2^, *P =* 0.003) and abaxial (inland ecotype effect = -63.13 mm^-2^, *P =* 0.004) leaf surfaces was greater in coastal than inland monkeyflower (Figure 6A, Appendices S11, S12, S13, S14), potentially offering more opportunities for saltwater entry in coastal leaves. Stomatal lengths were similar between ecotypes (adaxial inland effect = 0.996 µm, *P* = 0.25, abaxial inland effect = 0.62 µm, *P* = 0.40, Figure 6B, Appendices S11, S12, S13, S14). Thus, coastal plants allocated a higher estimated fraction of the epidermis area to stomata on both sides of the leaf (adaxial inland ecotype effect = -0.019, *P =* 0.015, abaxial inland effect = -0.013, *P* < 0.0001, Figure 6C, Appendices S11, S12, S13, S14). Stomatal conductance on the adaxial surface was similar between coastal and inland ecotypes (inland ecotype effect = 0.069 mol m^-2^ s^-1^, *P* = 0.28, Figure 6D, Appendices S12, S13); however, we note that replication was substantially less for stomatal conductance than the other traits we measured. These findings suggest that preventing saltwater entry by having few stomata, small stomata, or closed adaxial stomata are not salt spray exclusion strategies in coastal yellow monkeyflower.

**Figure 6.**
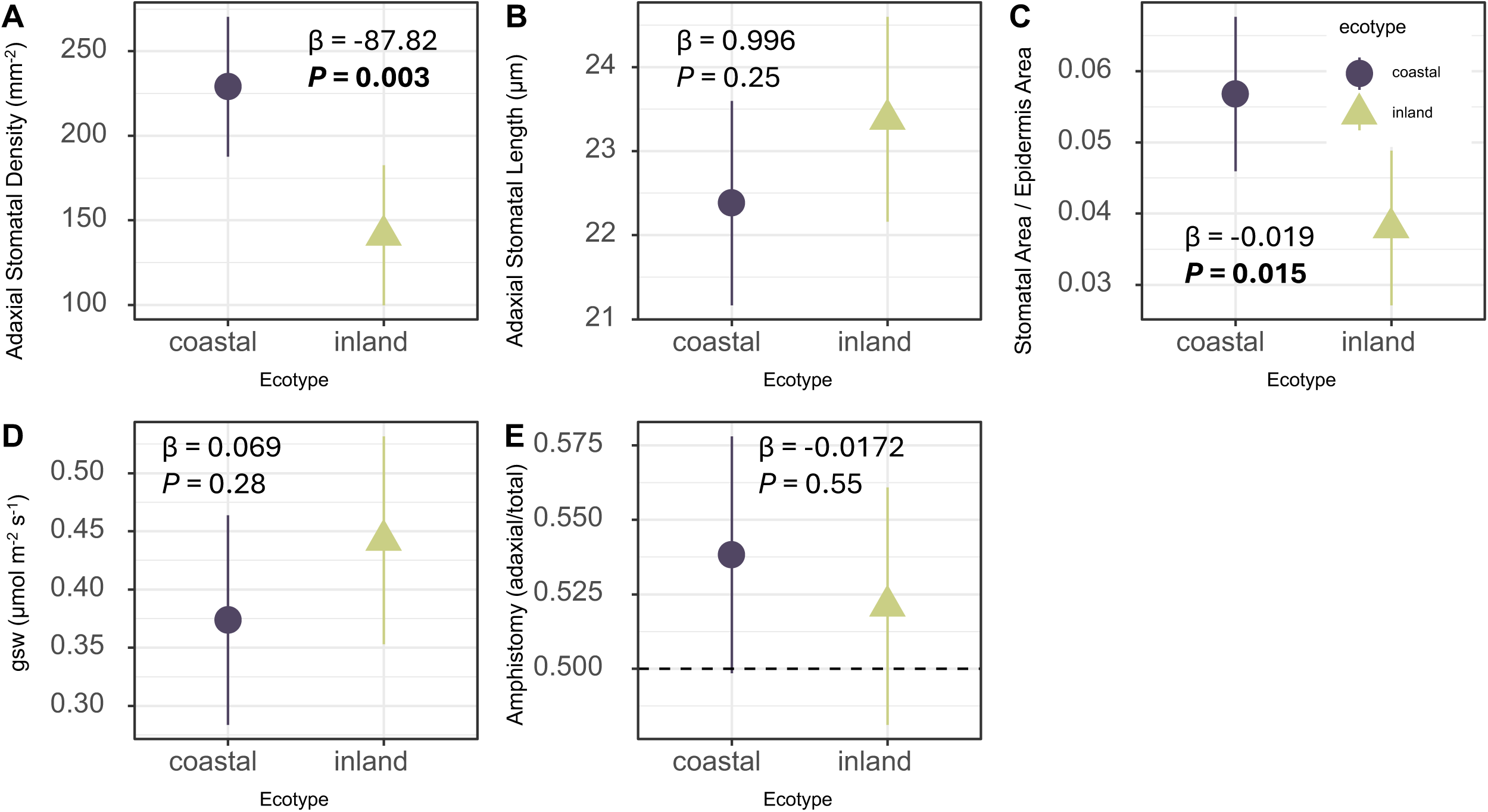
Comparison of adaxial stomatal traits between ecotypes. Points indicate LSMs for each ecotype from general linear mixed models, with 15 individuals measured per accession (except g_sw_ (D), where 6 leaves sampled from 2 – 3 individuals were measured per accession), and error bars show 95% confidence intervals. β and p-values indicate the effect and significance of inland ecotype on the trait. (A) Stomatal density in stomata per mm^2^ (y-axis) against ecotype (x-axis). (B) Stomatal length in µm (y-axis) against ecotype (x-axis). (C) Estimated fraction of epidermis area allocated to stomata (y-axis) against ecotype (x-axis). (D) Stomatal conductance in µmol m^-2^s^-1^ (y-axis) against ecotype (x-axis). (E) Amphistomy as adaxial stomatal count / abaxial stomatal count in the same size field of view (y-axis) against ecotype (x-axis). Dashed line indicates amphistomy = 0.5, which corresponds to equal numbers of stomata on the adaxial and abaxial surfaces.

Although most plants produce stomata primarily on the abaxial side of the leaf, the congener *Mimulus cardinalis* Dougl. ex Benth is amphistomatous (Muir, 2015; Nelson et al., 2021). We expected salt spray to land mainly on the more exposed adaxial surface, which may drive differences in stomatal traits on the two leaf sides (as seen for cuticular and wettability traits in *Festuca rubra* subsp. *litoralis* (G.Mey.) Auquier, Humphreys et al., 1986). However, amphistomy (the ratio of adaxial:total stomatal density) was similar between ecotypes (inland ecotype effect = -0.0172, *P* = 0.55, Figure 6E, Appendices S12, S13), indicating that stomatal placement on the abaxial side to prevent seawater entry is not a salt exclusion strategy in coastal yellow monkeyflower.

### Tissue tolerance

Coastal leaves were more succulent than inland leaves (inland ecotype effect = -0.0065 g/cm^2^, *P* = 0.012, Figure 7A, Appendix S15). This is consistent with greater tissue tolerance in the coastal ecotype, since succulence contributes to tissue tolerance by allowing sodium sequestration in large vacuoles and diluting sodium through high water content.

**Figure 7.**
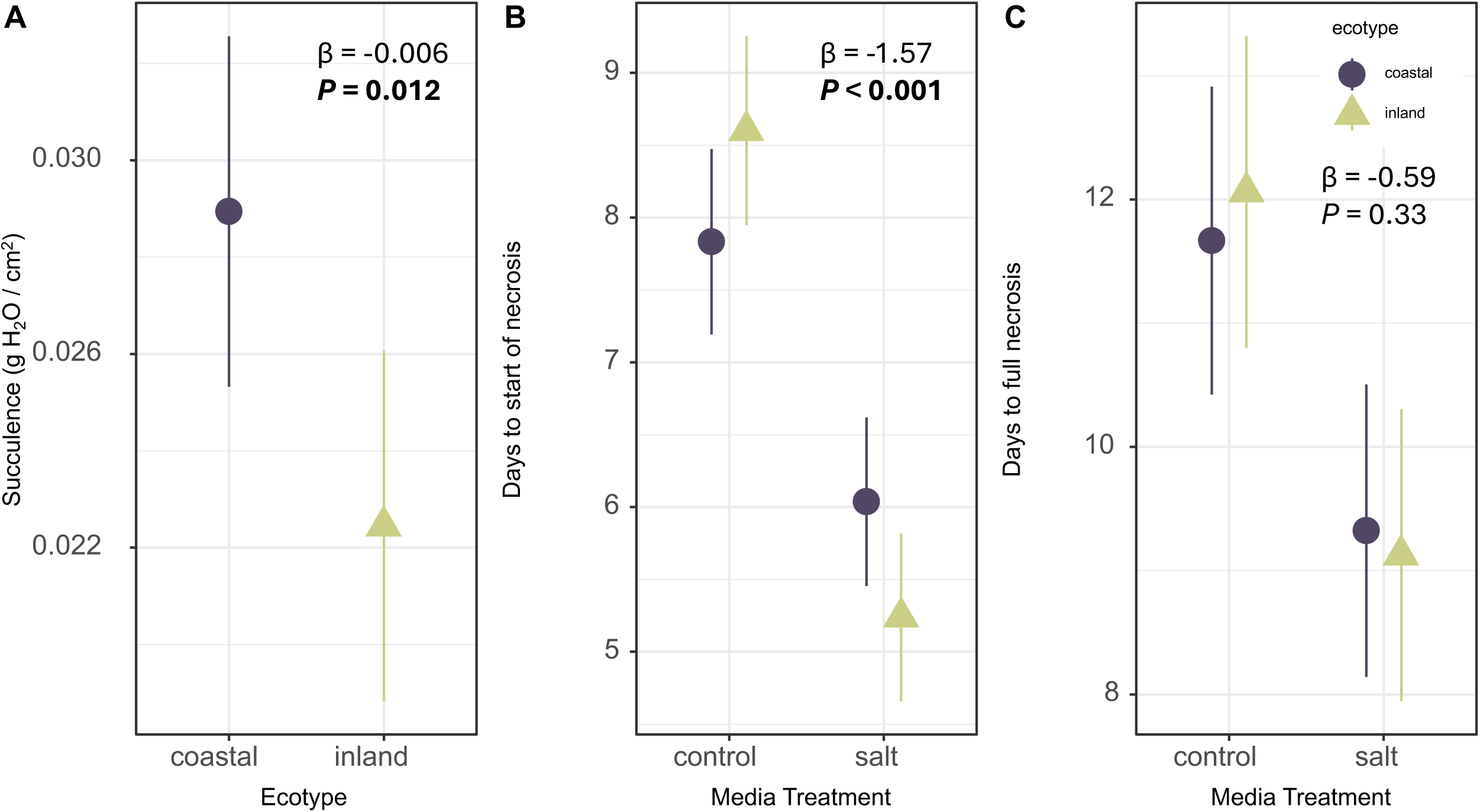
Comparison of leaf tissue tolerance traits between ecotypes. Error bars indicate 95% confidence intervals from general linear mixed models. (A) Succulence as (fresh mass - dry mass)/leaf area, in g H_2_O/cm^2^. β and p-value indicate the effect and significance of inland ecotype on the trait from a general linear mixed model. (B) Number of days to the start of visible tissue necrosis for leaf disks from coastal and inland accessions that were plated on media with 100 mM NaCl (salt) or no added NaCl (control). Points indicate LSMs for each ecotype and treatment combination, with 6–10 leaf disks scored for each accession under each treatment. β and p-value indicate the effect size and significance of the interaction between salt treatment and inland ecotype. (C) Number of days to complete tissue necrosis for leaf disks from coastal and inland accessions that were plated on media with 100 mM NaCl (salt) or no added NaCl (control). Points indicate LSMs for each ecotype and treatment combination, with 6–10 leaf disks scored for each accession under each treatment. β and p-value indicate the effect size and significance of the interaction between salt treatment and inland ecotype.

The more direct test for tissue tolerance differences, using leaf disks plated on NaCl media, revealed further evidence for higher leaf tissue tolerance of coastal than inland plants. Leaf disks from coastal and inland accessions took similar amounts of time to begin showing necrosis on control media, but inland accessions began showing necrosis faster than coastal accessions on salt media (treatment*ecotype interaction effect = -1.5674 days, *P* = 0.0001, Figure 7B, Appendix S16). However, the effect of treatment on time to full tissue necrosis did not differ between ecotypes (treatment*ecotype interaction effect = -0.59 days, *P* = 0.33, Figure 7C, Appendix S16).

## DISCUSSION

In this study, we found evidence that a complex complementary set of mechanisms contributes to salt spray adaptation in coastal perennial monkeyflower. We present the first evidence of salt spray exclusion in this species and find greater succulence in coastal perennials, which may contribute to observed differences in tissue tolerance.

Together, salt spray exclusion and succulence led to lower sodium concentrations in leaf tissue after salt spray than inland plants. Interestingly, salt spray led to increased LMA and decreased leaf succulence in the inland annual, which are not widely documented responses in the salinity stress literature. While salt concentrations reached higher levels in inland than coastal leaves, our efforts to understand this phenomenon through our investigation of multiple candidate leaf traits for salt spray exclusion (including stomatal traits and reduced leaf wettability) did not uncover the ultimate mechanisms of salt spray exclusion in this system. Beyond exclusion, we found compelling evidence for higher tissue tolerance in coastal plants than in inland plants through leaf disk assays (based on the time to start of necrosis, but not the time to full necrosis), which is potentially mediated by higher succulence in coastal leaves. Taken together, our study reveals that salt spray exclusion and tissue tolerance mechanisms contribute to the resilience of coastal monkeyflower to salt spray and suggests that local adaptation to oceanic salt spray in plants may often involve the combination of diverse developmental and physiological mechanisms (Figure 1).

Research on yellow monkeyflowers has now established that salt spray drives local adaptation to the coastal habitat, and tested three sequentially acting mechanisms of salt spray resilience. Salt spray, not soil salinity, is the major selective agent driving local adaptation to the coastal habitat (Popovic and Lowry, 2020). Salt spray carried by intense spring winds selects for short plants and late flowering in most coastal accessions (Toll et al., 2025), with the shortest coastal accessions found closest to the coast and in the windier northern part of the range (Zambiasi and Lowry, 2024). We found evidence for two additional mechanisms, salt spray exclusion and tissue tolerance, that occur after saltwater contacts the plant. Once saltwater lands on a leaf surface, coastal monkeyflowers prevent sodium ions from entering the leaf tissue (Figure 3B). Moreover, if saltwater does enter the leaf tissue, coastal leaves prevent necrosis longer than inland leaves, indicating greater tissue tolerance (Figure 7B, Appendix S15). Previous work showed that coastal plants survive longer than inland plants when grown in sodium chloride solution, suggesting that the coastal accession had greater tissue tolerance to sodium (Lowry et al., 2009). Our study expands on this previous work by showing that these differences in tissue tolerance (and salt spray exclusion) are ecotypic, replicated across multiple coastal and inland accessions.

Greater succulence in leaves of the coastal ecotype (Figure 7A) and less loss of succulence upon salt treatment (Figure 3C) may contribute to greater tissue tolerance in the coastal ecotype. Like many abiotic stressors, resilience under salt spray calls for multiple strategies that act at multiple stages: before salt reaches the plant, at the leaf surface, and inside plant cells.

While our research has now demonstrated that coastal monkeyflowers employ three salt spray resilience strategies (Figure 1), additional research is needed to identify the physiological mechanisms of salt spray exclusion and tissue tolerance, and the genetic mechanisms of all three strategies. Despite investigating stomatal density, size, and conductance, as well as leaf wettability traits, we were unable to identify a leaf surface trait that underlies observed salt spray exclusion in the coastal ecotype. All candidate leaf traits investigated for a potential role in salt spray exclusion were either similar between ecotypes (such as contact angle, Figure 5A-5B) or the difference was opposite of what would be required to facilitate salt spray exclusion in the coastal ecotype (such as stomatal density, Figure 6A, and leaf water drop adhesion assay, Figure 5C). Further experiments on the mechanism of salt spray exclusion should examine stomatal responses to salt spray, since stomatal closure in response to sodium may be an adaptive salt spray exclusion mechanism. We also note that our measurements of stomatal conductance had less replication than other parts of our study and examined only the adaxial side; a more thorough investigation of stomatal opening on both leaf surfaces under salt spray conditions may uncover a role in salt spray exclusion. Sodium ions cause stomatal opening in most plants, but rapid stomatal closure in the halophyte *Aster tripolium* L. upon leaf epidermis treatment with sodium chloride suggests that stomatal regulation in response to sodium may be a key part of adaptation to salinity stress (Perera et al., 1994). Investigating stomatal responses to salt spray and other unmeasured leaf traits that influence foliar water uptake and water shedding from leaves, such as trichome density, cuticular anatomy, and petiole rigidity (Berry et al., 2019; Lenz et al., 2022), as well as determining the path through which sodium enters leaves, may uncover the mechanism of salt spray exclusion.

Towards understanding the tissue tolerance mechanism, we showed that coastal monkeyflowers have higher tissue tolerance than inland monkeyflowers (Figure 7B). While leaf disks are far-removed from a whole plant surviving in nature, they allowed us to examine the role of tissue tolerance alone after eliminating the salt spray escape and exclusion strategies that contribute to survival in the field. Our finding of greater tissue tolerance in coastal monkeyflowers is consistent with a previous finding that a coastal accession had greater survival when salt entered the plant through a hydroponic solution than an inland accession, in a context where salt exclusion at the leaf surface is not possible (Lowry et al., 2009). Our study expands this finding of greater tissue tolerance in coastal than inland monkeyflowers to multiple accessions across a broad geographic range, indicating that tissue tolerance to sodium is a widespread characteristic of the coastal ecotype. Additional research should investigate whether the greater succulence observed in this study in coastal than inland monkeyflower (Figure 7A, 3C) *contributes* to their greater tissue tolerance, for example, by allowing plants to sequester sodium in large vacuoles (Apse et al., 2003). Such studies would offer a more complete understanding of how the multiple salt spray resilience strategies work on physiological and molecular levels.

Additional untested mechanisms of salt spray resilience may also exist. When plant roots are exposed to salt, the sodium ions outside the root can cause osmotic stress and loss of turgor when water is drawn out (Munns & Tester, 2008). A similar mechanism in leaves could explain our finding of water loss upon salt spray (Figure 3C). Coastal plants could be less susceptible to water loss caused by the osmotic pressure of sodium chloride encrusting the leaf surface, or could more readily replenish lost water from the soil. In smooth cordgrass (*Spartina alterniflora* Loisel.), osmotic potential decreased when plants were treated with salt spray (Touchette *et al*., 2009). In the orchid *Epidendrum fulgens* Brongn., salt spray treatment reduced the rate of leaf expansion, which may indicate osmotic stress, but relative water content and osmotic potential were not affected by salt spray treatment (de Lima *et al*., 2024). To our knowledge, this is the first report of water loss due to salt spray on leaves, with other studies reporting *increased* succulence with salt spray treatment that was generally monitored over longer timescales (Boyce, 1954; De Vos et al., 2010). We speculate that the increase in succulence generally observed longer timescales reflects an increase in tissue tolerance by diluting sodium and/or producing larger vacuoles for sodium sequestration (Ogburn and Edwards, 2010; Munns et al., 2016). Our finding of a loss of succulence within hours of salt spray, particularly in the inland annual, suggests that the immediate response to salt spray may hinder these tissue tolerance mechanisms. The impact of salt spray on plant water relations in nature is likely influenced by coastal fog, salt spray intensity, and the length of time plants are exposed to salt spray, with this interplay potentially contributing to mortality differences between coastal and inland ecotypes under salt spray.

Another untested mechanism, salt secretion, may reduce sodium accumulation per unit leaf area, instead of or in addition to salt exclusion. Although yellow monkeyflowers lack conspicuous secretory structures such as the epidermal salt bladders in *Chenopodium quinoa* Willd. (Kiani-Pouya et al., 2017), some salt secretion may occur without specialized salt secretory structures. Salt may be secreted by glandular trichomes as in some halophytes (Flowers et al., 1990). We also observe substantial guttation on plants in the early morning, which may flush salt out of the leaves, although evidence that this secretion pathway substantially reduces shoot sodium levels in other systems is limited (Hossain et al., 2016).

The leaf traits measured in our study affect plant physiology beyond their potential role in salt spray resilience, which may influence resilience to other abiotic stressors. For example, water loss and carbon dioxide uptake both occur through stomata, and stomatal morphology can influence not only the entry of salt water but also plant water relations and photosynthesis (Bertolino et al., 2019). Further, amphistomy, the placement of stomata on both sides of a leaf, allows CO_2_ to diffuse from stomatal openings to chloroplasts more rapidly during photosynthesis (Muir, 2015; Triplett et al., 2024) but can also increase the risk of taking up saltwater through exposed adaxial stomata. And while leaf wettability affects salt spray entry (Ahmad and Wainwright, 1976), it also has other ecophysiological consequences (reviewed in Lenz et al., 2022). Wet leaves can access supplemental water from fog, dew, or canopy rainwater (Steppe et al., 2018; Cavallaro et al., 2020, 2022), face slower CO_2_ diffusion and reduced photosynthesis (Brewer and Smith, 1995), and are more susceptible to fungal pathogens (Bradley et al., 2003) than dry leaves. In yellow monkeyflower, the coastal and inland populations face numerous abiotic stressors that drive local adaptation, including herbivory in coastal habitats and terminal summer drought in inland habitats (Hall and Willis, 2006; Lowry et al., 2019a; Popovic and Lowry, 2020). Some leaf traits examined for their potential influence on salt spray resilience, such as amphistomy and leaf wettability, may be primarily driven by selection for photosynthetic rate, drought resilience, or other factors.

In addition to the complex interplay between salt spray and other stressors, plants displaying salt spray resilience mechanisms may incur an energetic cost. When soil salinity-tolerant plants sequester sodium in the vacuole, they establish an increased H^+^ concentration in the vacuole compared to the cytosol, allowing Na^+^/H^+^ antiporters (NHX) to move sodium from the cytosol to the vacuole (Rea and Sanders, 1987; Apse et al., 2003). Since establishing the H^+^ gradient requires ATP hydrolysis, vacuolar sequestration can be energetically costly (Shabala et al., 2020). Some plants maintain homeostasis under osmotic stress from saline soils by osmotic adjustment with organic solutes, which is also resource-intensive (Munns et al., 2020). Coastal perennials display a slower, more resource-conservative life history strategy than inland annuals, investing in stress tolerance mechanisms and defense rather than rapid growth (Lowry et al., 2019a) and may therefore dedicate resources to energetically expensive salt resilience mechanisms. Moreover, the need for inland annuals to complete their life cycle before a summer terminal drought may preclude them from dedicating water resources towards succulent leaves like the coastal perennials.

Just as in a prior study in this system (Wu et al., 2010), we observed greater LMA in coastal perennials than inland annuals, which is consistent with the slower, resource-conservative strategy of coastal perennials. Surprisingly, we found that salt spray led to an increase of LMA just 18 hours after salt spray treatment, with a particularly dramatic increase in inland annuals (Figure 4). Assuming equimolar amounts of sodium and chloride uptake in these leaves, the measured sodium is not sufficient to account for increased LMA based on sodium chloride alone. We speculate that salt-sprayed leaves may be mobilizing osmolytes from other parts of the plant towards the salt-sprayed leaves, or that chloride uptake is greater than sodium uptake in inland annuals, to give the observed increase in LMA under salt spray. LMA also increased under salinity treatment over longer timescales in both emergent and free-floating *Alternanthera philoxeroides* (Grewell et al., 2025). Comparing morphology, resource allocation, and stress responses of coastal perennials with inland perennials (e.g. Toll et al., 2025) could disentangle the roles of life history differences from salt spray tolerance. Exploring the energetic cost of salt spray exclusion and characterizing the salt spray exclusion and tissue tolerance mechanisms in yellow monkeyflowers would further our understanding of selection pressures on coastal-inland gradients.

Looking forward, changing wind patterns, increased wave action, and sea level rise under climate change are expected to cause increased plant exposure to salt spray (Young and Ribal, 2019; Du and Hesp, 2020). Like other stresses related to climate change, sensitive plant populations will respond to novel or increased salt spray by acclimating, adapting, shifting their range, or becoming locally extinct (Corlett and Westcott, 2013; Halpin-McCormick et al., 2025). Understanding the mechanisms of salt spray adaptation may allow us to uncover useful genetic variation, identify and conserve vulnerable plant species, and retain the ecological services provided by coastal plant communities.

## Acknowledgements

This work was supported by a National Science Foundation Graduate Research Fellowship to MLP (DGE-184-8739) and a National Science Foundation Division of Integrative Organismal Systems Grant to DBL (IOS-2153100). Ritta Mouayed, Jeffrey Yang, Elle Mader, and Arjun Kuruppu Goonetilleke assisted with image analysis. We thank V Pargulski and Berkley Walker for training with the LI-600. We thank Hui-Kyong Cho and Ilyeong Choi for training and assistance with MP-AES. We thank Nate Emery, Milagros del Pilar Jimenez-Hernandez, and Thomas Zambiasi for performing preliminary salt spray experiments. We thank the Big Sur Land Trust for permission to collect the TOR accession, and the State Parks of California and Oregon for additional collection permissions. We thank Jeffrey Conner, Lauren Sullivan, members of the Lowry lab, and two anonymous peer reviewers for helpful comments on the manuscript and analyses.

## Author Contributions

MLP, PCD, and KE designed and performed experiments and analyzed data. DBL conceived of the study and secured funding. MLP wrote the original manuscript draft. All authors read and approved the final manuscript.

## Data Availability

Data, metadata, and R scripts used in this study are available at the following GitHub repository: https://github.com/plunkert/Salt-spray-adaptation-2025 and archived on Zenodo (DOI 10.5281/zenodo.20936681): https://zenodo.org/records/20936681

## Supporting Information

Additional Supporting Information may be found online in the supporting information section at the end of the article.

**Appendix S1.** (A) Coastal perennial ecotype in coastal habitat (B) Inland annual ecotype in inland habitat.

**Appendix S2.** Coordinates for accessions used in this study.

**Appendix S3.** Standard curve for sodium quantification by MP-AES analysis.

**Appendix S4.** Methodological verification of rinsing step in leaf sample preparation for MP-AES analysis.

**Appendix S5.** Analyses of sodium concentration, salt exclusion, and succulence under salt treatment.

**Appendix S6.** Sodium and water levels in coastal and inland accessions.

**Appendix S7.** Leaf mass per area (y-axis) in coastal and inland accessions after salt spray or control treatment.

**Appendix S8.** Analysis of LMA (g m^-2^) under salt treatment.

**Appendix S9.** Wettability traits (y-axis) in coastal and inland accessions.

**Appendix S10.** Analyses of leaf wettability in coastal and inland accessions.

**Appendix S11.** Comparison of abaxial stomatal traits between ecotypes.

**Appendix S12.** Stomatal traits compared between coastal and inland accessions.

**Appendix S13.** Analyses of stomatal traits in coastal and inland accessions.

**Appendix S14.** Analyses of abaxial stomatal traits in coastal and inland accessions.

**Appendix S15.** Analysis of succulence in coastal and inland accessions.

**Appendix S16.** Analysis of tissue tolerance assay.

### Box 1. Proposed mechanisms of salt spray resilience.

**Salt spray escape** - features of plant architecture or phenology that reduce exposure to salt spray (Figure 1A). Examples: prostrate growth, delayed flowering

**Salt spray exclusion** - once salt spray lands on a plant, surface characteristics prevent it from entering tissues (Figure 1B). Examples: few or small stomata, reduced wettability through cuticle properties

**Tissue tolerance** - once salt enters plant tissues, tissue- and/or cell-level mechanisms prevent damage and maintain homeostasis (Figure 1C). Examples: reactive oxygen species (ROS) scavenging, sequestering sodium in large vacuoles, high tissue water content preventing damaging sodium concentrations

## REFERENCES

Ahmad, I., and S. J. Wainwright. 1976. Ecotype differences in leaf surface properties of *Agrostis stolonifera* from salt marsh, spray zone, and inland habitats. New Phytologist 76: 361–366.

Apse, M. P., J. B. Sottosanto, and E. Blumwald. 2003. Vacuolar cation/H+ exchange, ion homeostasis, and leaf development are altered in a T-DNA insertional mutant of AtNHX1, the *Arabidopsis* vacuolar Na+/H+ antiporter. The Plant Journal 36: 229–239.

Barbour, M. G. 1978. Salt spray as a microenvironmental factor in the distribution of beach plants at Point Reyes, California. Oecologia 32: 213–224.

Barker, W. R., G. Nesom, P. M. Beardsley, and N. S. Fraga. 2012. A taxonomic conspectus of Phrymaceae: A narrowed circumscription for *Mimulus*, new and resurrected genera, and new names and combinations. Phytoneuron 39: 1–60.

Berry, Z. C., N. C. Emery, S. G. Gotsch, and G. R. Goldsmith. 2019. Foliar water uptake: Processes, pathways, and integration into plant water budgets. *Plant*, Cell & Environment 42: 410–423.

Bertolino, L. T., R. S. Caine, and J. E. Gray. 2019. Impact of stomatal density and morphology on water-use efficiency in a changing world. Frontiers in Plant Science 10: 225.

Boyce, S. G. 1954. The salt spray community. Ecological Monographs 24: 29–67.

Bradley, D. J., G. S. Gilbert, and I. M. Parker. 2003. Susceptibility of clover species to fungal infection: the interaction of leaf surface traits and environment. American Journal of Botany 90: 857–864.

Brewer, C. A., and W. K. Smith. 1995. Leaf surface wetness and gas exchange in the pond lily *Nuphar polysepalum* (Nymphaeaceae). American Journal of Botany 82: 1271–1277.

Brooks, M., K. Kristensen, K. van Benthem, A. Magnusson, C. Berg, A. Nielsen, H. Skaug, et al. 2017. glmmTMB balances speed and flexibility among packages for zero-inflated generalized linear mixed modeling. The R Journal 9: 378–400.

Brugnara, M. 2006. Contact angle. ImageJ Plugins. Website https://imagej.net/ij/plugins/contact-angle.html [accessed 26 June 2025].

Burkhardt, J., S. Basi, S. Pariyar, and M. Hunsche. 2012. Stomatal penetration by aqueous solutions – an update involving leaf surface particles. New Phytologist 196: 774–787.

Cavallaro, A., L. Carbonell-Silletta, A. Burek, G. Goldstein, F. G. Scholz, and S. J. Bucci. 2022. Leaf surface traits contributing to wettability, water interception and uptake of above-ground water sources in shrubs of Patagonian arid ecosystems. Annals of Botany 130: 409–418.

Cavallaro, A., L. Carbonell Silleta, D. A. Pereyra, G. Goldstein, F. G. Scholz, and S. J. Bucci. 2020. Foliar water uptake in arid ecosystems: seasonal variability and ecophysiological consequences. Oecologia 193: 337–348.

Clausen, J., and W. M. Hiesey. 1958. Genetic structure of ecological races. The Lord Baltimore Press, Inc, Baltimore, Maryland, USA.

Corlett, R. T., and D. A. Westcott. 2013. Will plant movements keep up with climate change? Trends in Ecology & Evolution 28: 482–488.

Coughlan, J. M., M. W. Brown, and J. H. Willis. 2021. The genetic architecture and evolution of life-history divergence among perennials in the *Mimulus guttatus* species complex. Proceedings of the Royal Society B: Biological Sciences 288: 20210077.

De Vos, A. C., R. Broekman, M. P. Groot, and J. Rozema. 2010. Ecophysiological response of *Crambe maritima* to airborne and soil-borne salinity. Annals of Botany 105: 925–937.

Du, J., and P. A. Hesp. 2020. Salt spray distribution and its impact on vegetation zonation on coastal dunes: a review. Estuaries and Coasts 43: 1885–1907.

Emery, N. C. 2016. Foliar uptake of fog in coastal California shrub species. Oecologia 182: 731–742.

Flowers, T. J., S. A. Flowers, M. A. Hajibagheri, and A. R. Yeo. 1990. Salt tolerance in the halophytic wild rice, *Porteresia coarctata* Tateoka. New Phytologist 114: 675–684.

Frayer, M. E., H. K. Soliman, P. F. Schwarz, and J. M. Coughlan. 2026. Introgression and parental conflict shape repeated occurrences of postzygotic isolation in *Mimulus*. Current Biology 36: 1454–1467.e4.

Gould, B. A., Y. Chen, and D. B. Lowry. 2017. Pooled ecotype sequencing reveals candidate genetic mechanisms for adaptive differentiation and reproductive isolation. Molecular Ecology 26: 163–177.

Grewell, B. J., B. Gallego-Tévar, J. M. Castillo, C. J. Futrell, R. E. Drenovsky, N. E. Harms, and P. D. Pratt. 2025. Functional trait responses of emergent and free-floating *Alternanthera philoxeroides* to increasing salinity with sea level rise: stress tolerance, avoidance, and escape strategies. NeoBiota 102: 9–36.

Grieve, A. M., and M. G. Pitman. 1978. Salinity damage to Norfolk island pines caused by surfactants. III. Evidence for stomatal penetration as the pathway of salt entry to leaves. Functional Plant Biology 5: 397–413.

Guo, B., S. K. Arndt, R. E. Miller, C. Szota, and C. Farrell. 2023. How does leaf succulence relate to plant drought resistance in woody shrubs? Tree Physiology 43: 1501–1513.

Guzmán-Delgado, P., E. Laca, and M. A. Zwieniecki. 2021. Unravelling foliar water uptake pathways: The contribution of stomata and the cuticle. *Plant*, Cell & Environment 44: 1728–1740.

Hall, M. C., and J. H. Willis. 2006. Divergent selection on flowering time contributes to local adaptation in *Mimulus guttatus* populations. Evolution 60: 2466–2477.

Halpin-McCormick, A., T. Sherrill, C. Davenport, D. Wolkis, S. K. Walsh, and K. E. Barton. 2025. Sensitivity to elevated salinity in coastal dune plants. Oecologia 207: 1–12.

Haq, T. U., J. Akhtar, K. A. Steele, R. Munns, and J. Gorham. 2013. Reliability of ion accumulation and growth components for selecting salt tolerant lines in large populations of rice. Functional Plant Biology 41: 379–390.

Hossain, Md. B., N. Matsuyama, and M. Kawasaki. 2016. Hydathode morphology and role of guttation in excreting sodium at different concentrations of sodium chloride in eddo. Plant Production Science 19: 528–539.

Humphreys, M. O., M. P. Kraus, and R. G. Wyn Jones. 1986. Leaf-surface properties in relation to tolerance of salt spray in *Festuca rubra* ssp. *litoralis* (G. F. W. Meyer) Aquier. New Phytologist 103: 717–723.

Itoh, M. 2021. Phenotypic variation and adaptation in morphology and salt spray tolerance in coastal and inland populations of *Setaria viridis* in central Japan. Weed Research 61: 199–209.

Itoh, M., K. Fukunaga, and T. Osako. 2024. Local adaptation in parapatric and sympatric mosaic coastal habitats through trait divergence of *Setaria viridis*. Journal of Ecology 112: 784–799.

Kiani-Pouya, A., U. Roessner, N. S. Jayasinghe, A. Lutz, T. Rupasinghe, N. Bazihizina, J. Bohm, et al. 2017. Epidermal bladder cells confer salinity stress tolerance in the halophyte quinoa and *Atriplex* species. *Plant*, Cell & Environment 40: 1900–1915.

Lenth, R. V., B. Banfai, B. Bolker, P. Buerkner, I. Giné-Vázquez, M. Herve, M. Jung, et al. 2025. emmeans: estimated marginal means, aka least-squares means. Website https://cran.r-project.org/web/packages/emmeans/index.html [accessed 24 July 2025].

Lenz, A.-K., U. Bauer, and G. D. Ruxton. 2022. An ecological perspective on water shedding from leaves. Journal of Experimental Botany 73: 1176–1189.

Liu, R., T. Wang, Q. Li, L. Wang, and J. Song. 2024. The role of tissue succulence in plant salt tolerance: an overview. Plant Growth Regulation 103: 283–292.

Lovell, J. T., R. Walstead, A. Lawrence, E. Stark-Dykema, M. C. Farnitano, A. Harder, T. Brůna, et al. 2025. Comparative analyses of four reference genomes reveal exceptional diversity and weak linked selection in the yellow monkeyflower (*Mimulus guttatus*) complex. Molecular Ecology Resources 25: e70012.

Lowry, D. B., D. Popovic, D. J. Brennan, and L. M. Holeski. 2019a. Mechanisms of a locally adaptive shift in allocation among growth, reproduction, and herbivore resistance in *Mimulus guttatus*. Evolution 73: 1168–1181.

Lowry, D. B., and J. H. Willis. 2010. A widespread chromosomal inversion polymorphism contributes to a major life-history transition, local adaptation, and reproductive isolation. PLOS Biology 8: e1000500.

Lowry, D. B., J. M. Sobel, A. L. Angert, T.-L. Ashman, R. L. Baker, B. K. Blackman, Y. Brandvain, et al. 2019b. The case for the continued use of the genus name *Mimulus* for all monkeyflowers. Taxon 68: 617–623.

Lowry, D. B., M. C. Hall, D. E. Salt, and J. H. Willis. 2009. Genetic and physiological basis of adaptive salt tolerance divergence between coastal and inland *Mimulus guttatus*. New Phytologist 183: 776–788.

Lowry, D. B., R. C. Rockwood, and J. H. Willis. 2008. Ecological reproductive isolation of coast and inland races of *Mimulus guttatus*. Evolution 62: 2196–2214.

Muir, C. D. 2015. Making pore choices: repeated regime shifts in stomatal ratio. Proceedings of the Royal Society B: Biological Sciences 282: 20151498.

Muir, C. D., M. À. Conesa, J. Galmés, V. S. Pathare, P. Rivera, R. López Rodríguez, T. Terrazas, and D. Xiong. 2023. How important are functional and developmental constraints on phenotypic evolution? An empirical test with the stomatal anatomy of flowering plants. The American Naturalist 201: 794–812.

Munns, R., J. B. Passioura, T. D. Colmer, and C. S. Byrt. 2020. Osmotic adjustment and energy limitations to plant growth in saline soil. New Phytologist 225: 1091–1096.

Munns, R., and M. Tester. 2008. Mechanisms of salinity tolerance. Annual Review of Plant Biology 59: 651–681.

Munns, R., R. A. James, M. Gilliham, T. J. Flowers, and T. D. Colmer. 2016. Tissue tolerance: an essential but elusive trait for salt-tolerant crops. Functional Plant Biology 43: 1103–1113.

Nelson, T. C., C. D. Muir, A. M. Stathos, D. D. Vanderpool, K. Anderson, A. L. Angert, and L. Fishman. 2021. Quantitative trait locus mapping reveals an independent genetic basis for joint divergence in leaf function, life-history, and floral traits between scarlet monkeyflower (*Mimulus cardinalis*) populations. American Journal of Botany 108: 844–856.

Nesom, G. L. 2014. Updated classifcation and hypothetical phylogeny of *Erythranthe* sect. Simiola (Phrymaceae). Phytoneuron 81: 1–6.

Ogburn, R. M., and E. J. Edwards. 2010. The ecological water-use strategies of succulent plants. In J.-C. Kader, and M. Delseny [eds.], Advances in botanical research, vol. 55, 179–225. Academic Press, London, UK.

Oneal, E., D. B. Lowry, K. M. Wright, Z. Zhu, and J. H. Willis. 2014. Divergent population structure and climate associations of a chromosomal inversion polymorphism across the *Mimulus guttatus* species complex. Molecular Ecology 23: 2844–2860.

Perera, L. K. R. R., T. A. Mansfield, and A. J. C. Malloch. 1994. Stomatal responses to sodium ions in *Aster tripolium*: a new hypothesis to explain salinity regulation in above-ground tissues. Plant, Cell & Environment 17: 335–340.

Popovic, D., and D. B. Lowry. 2020. Contrasting environmental factors drive local adaptation at opposite ends of an environmental gradient in the yellow monkeyflower (*Mimulus guttatus*). American Journal of Botany 107: 298–307.

R Core Team. 2024. R: a language and environment for statistical computing (Version 4.4.1 Software). R Foundation for Statistical Computing, Vienna, Austria.

Rajendran, K., M. Tester, and S. J. Roy. 2009. Quantifying the three main components of salinity tolerance in cereals. Plant, Cell & Environment 32: 237–249.

Rea, P. A., and D. Sanders. 1987. Tonoplast energization: Two H+ pumps, one membrane. Physiologia Plantarum 71: 131–141.

Schneider, C. A., W. S. Rasband, and K. W. Eliceiri. 2012. NIH Image to ImageJ: 25 years of image analysis. Nature Methods 9: 671–675.

Shabala, S., G. Chen, Z.-H. Chen, and I. Pottosin. 2020. The energy cost of the tonoplast futile sodium leak. New Phytologist 225: 1105–1110.

Steppe, K., M. W. Vandegehuchte, B. A. E. Van de Wal, P. Hoste, A. Guyot, C. E. Lovelock, and D. A. Lockington. 2018. Direct uptake of canopy rainwater causes turgor-driven growth spurts in the mangrove *Avicennia marina*. Tree Physiology 38: 979–991.

Toll, K., C. Crockett, L. Tocci, and D. B. Lowry. 2025. Seasonal variation in wind speed and oceanic salt spray favors delayed reproduction in coastal yellow monkeyflowers. 2025.11.07.687229.

Toll, K., M. Blanchard, A. Scharnagl, D. Lowry, and L. Holeski. 2024. Plant hormone manipulation impacts salt spray tolerance, which preempts herbivory as a driver of local adaptation in the yellow monkeyflower, Mimulus guttatus. 2024.05.23.595619.

Triplett, G., T. N. Buckley, and C. D. Muir. 2024. Amphistomy increases leaf photosynthesis more in coastal than montane plants of Hawaiian ilima (*Sida fallax*). American Journal of Botany 111: e16284.

Turesson, G. 1922. The genotypical response of the plant species to the habitat. Hereditas 3: 211–350.

Twyford, A. D., and J. Friedman. 2015. Adaptive divergence in the monkey flower *Mimulus guttatus* is maintained by a chromosomal inversion. Evolution 69: 1476–1486.

Wang, H., H. Shi, Y. Li, and Y. Wang. 2014. The effects of leaf roughness, surface free energy and work of adhesion on leaf water drop adhesion. PLOS ONE 9: e107062.

Wilkinson, M. J., F. Roda, G. M. Walter, M. E. James, R. Nipper, J. Walsh, S. L. Allen, et al. 2021. Adaptive divergence in shoot gravitropism creates hybrid sterility in an Australian wildflower. Proceedings of the National Academy of Sciences 118: e2004901118.

Willis, J. H. 1993. Partial self-fertilization and inbreeding depression in two populations of Mimulus guttatus. Heredity 71: 145–154.

Wu, C. A., D. B. Lowry, L. I. Nutter, and J. H. Willis. 2010. Natural variation for drought-response traits in the *Mimulus guttatus* species complex. Oecologia 162: 23–33.

Young, I. R., and A. Ribal. 2019. Multiplatform evaluation of global trends in wind speed and wave height. Science 364: 548–552.

Zambiasi, T., and D. B. Lowry. 2024. Ocean exposure and latitude drive multiple clines within the coastal perennial ecotype of the yellow monkeyflower, *Mimulus guttatus*. American Journal of Botany 111: e16402.

