## Appendix S1 for "Foliar salt spray exclusion and tissue tolerance underlie local adaptation to oceanic salt spray"

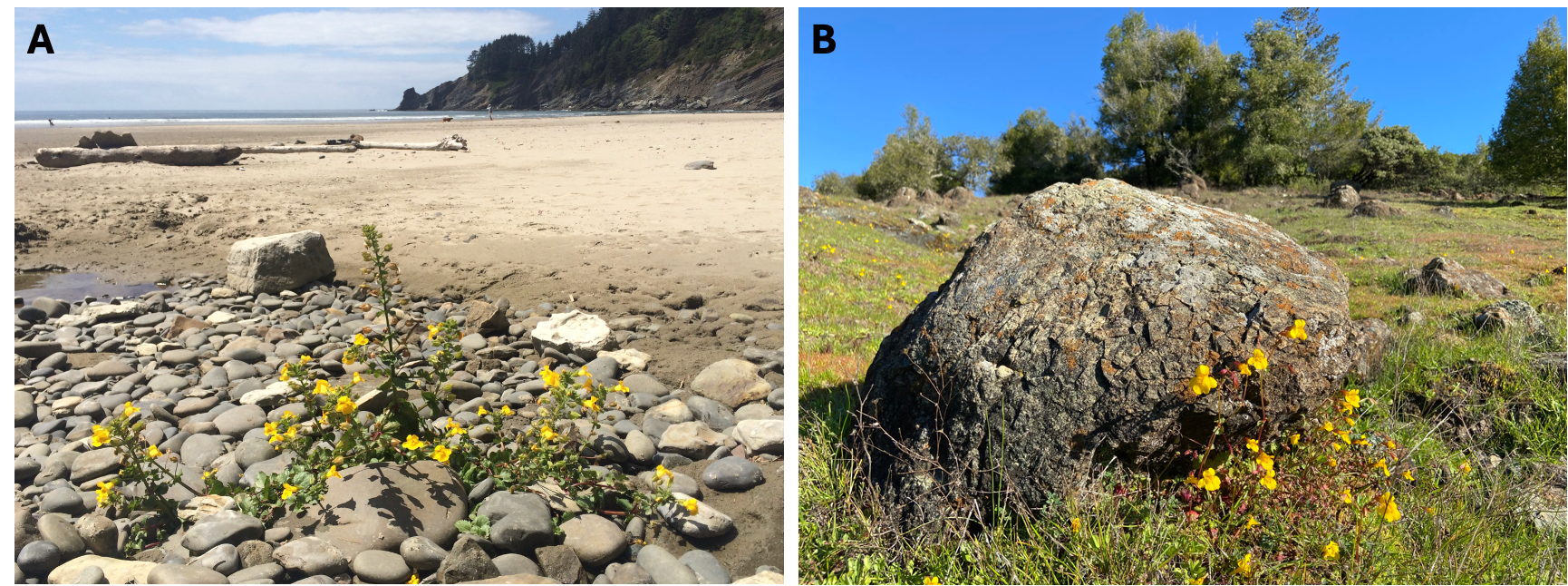

**Appendix S1.** (A) Coastal perennial ecotype in coastal habitat (OSW, Oswald West State Park, Oregon, USA, 45.761083°N, 123.966567°W). (B) Inland annual ecotype in inland habitat (MOR, Alder Creek Ranch on Morelli Rd, Occidental, California, USA, 38.4297°N, 122.951617°W).
