## Appendix S2 for "Foliar salt spray exclusion and tissue tolerance underlie local adaptation to oceanic salt spray"

**Appendix S2.** Coordinates for accessions used in this study.

| Population Code | Population Name | Family ID | Generations Inbred | Latitude | Longitude | Ecotype |
| --- | --- | --- | --- | --- | --- | --- |
| BHE | Bodega Headlands | 18-1 | 1 | 38.307033 | -123.05842 | coastal |
| HEC | Heceta Beach | 30-5 | 4 | 44.135067 | -124.1228 | coastal |
| LMC | Lower Mendocino County | 24 | 2 | 38.863983 | -123.08392 | inland |
| OAE | Occidental Arts and Ecology Center | 19 | 1 | 38.411267 | -122.95972 | inland |
| OPB | Otter Point State Park beach | 31-2 | 3 | 42.464017 | -124.42292 | coastal |
| PGR | Pacific Grove | 3 | 3 | 36.6279917 | -121.92083 | coastal |
| RGR | Rogue River | 27 | 1 | 42.48925 | -124.2084 | inland |
| SWB | Sperm Whale Beach | 4 | 2 | 39.035983 | -123.69047 | coastal |
| SWC | Sweet Creek Road | 24-9-2 | 7 | 43.959467 | -123.90245 | inland |
| TOR | Toro County Park | 3 | 2 | 36.603042 | -121.68841 | inland |
