## Supplementary figures and images for "Foliar salt spray exclusion and tissue tolerance underlie local adaptation to oceanic salt spray"

### Appendix S3

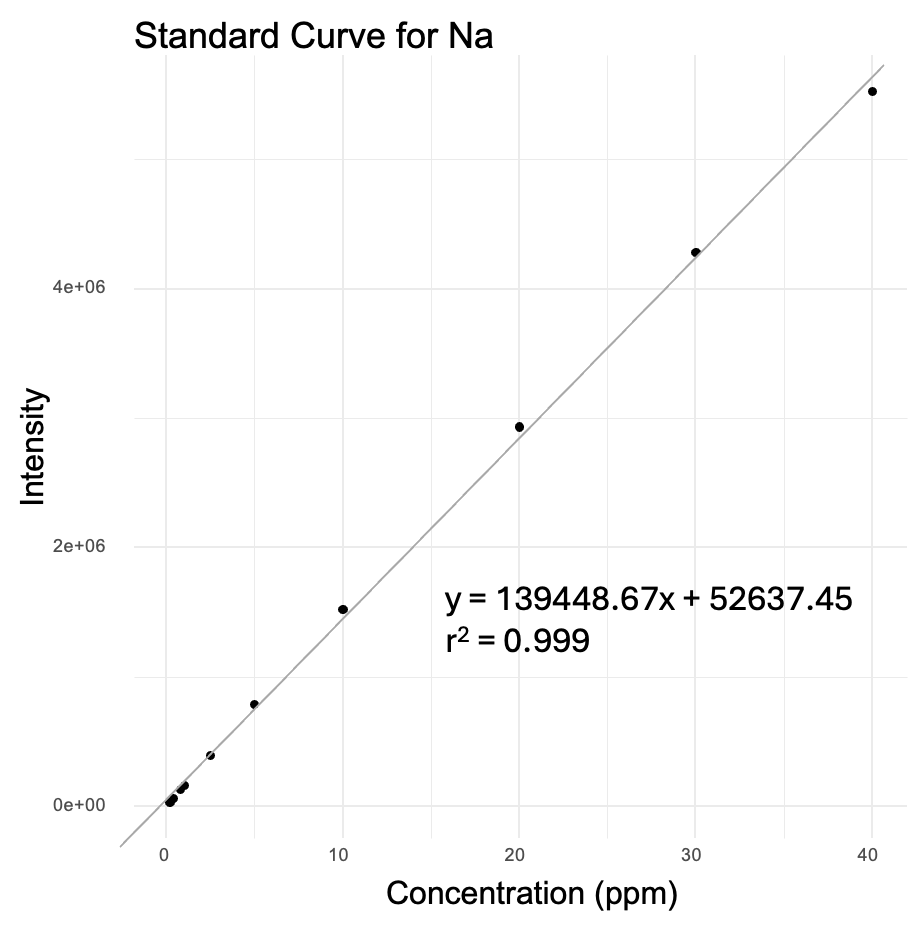

**Appendix S3.** Standard curve for sodium quantification by MP-AES analysis.
