## Appendix S4 for "Foliar salt spray exclusion and tissue tolerance underlie local adaptation to oceanic salt spray"

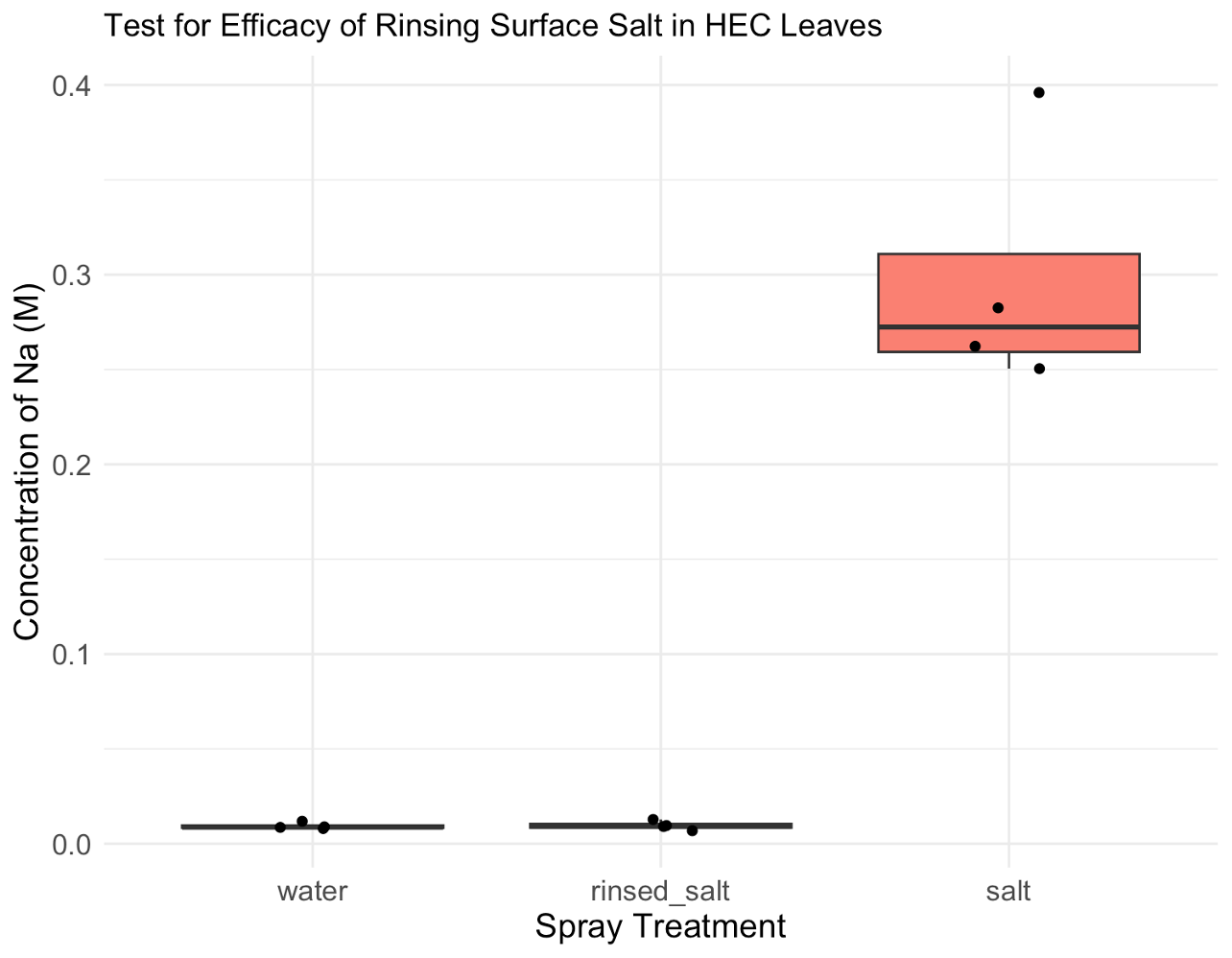

**Appendix S4.** Methodological verification of rinsing step in leaf sample preparation for MP-AES analysis. All leaves analyzed are from the HEC accession, with each point showing concentration of Na^+^ in the leaf calculated from MP-AES ppm. Water and salt treatments are identical to the water and salt treatments in the larger analysis. For the rinsed_salt treatment, the leaves were rinsed immediately after spraying salt without allowing time for sodium to enter the leaf tissue, so any increase in sodium concentration compared to the water treatment would reflect residual sodium left behind by poor rinsing. This test indicates that sodium measurements by MP-AES on rinsed leaves reflect sodium inside the leaf, not residual sodium on the leaf surface.
