## Appendix S5 for "Foliar salt spray exclusion and tissue tolerance underlie local adaptation to oceanic salt spray"

**Appendix S5.** Analyses of sodium concentration, salt exclusion, and succulence under salt treatment. General linear mixed models of sodium concentration (M), salt exclusion (as µmol of Na per cm^2^ of leaf area), and succulence (as g H_2_O per cm^2^ of leaf area). Effect sizes are shown in the units of the response variable.

Sodium concentration ~ ecotype * treatment + (1 | accession)

| Effect | Estimate | SE | *z*-value | *P*-value |
| --- | --- | --- | --- | --- |
| Intercept(coastal water) | 0.009 | 0.0264 | 0.34 | 0.73 |
| Ecotype(inland) | -0.001 | 0.0373 | -0.03 | 0.98 |
| Treatment(salt) | 0.1521 | 0.0368 | 4.13 | <0.0001 |
| Treatment(salt):Ecotype(inland) | 0.2917 | 0.0521 | 5.60 | <0.0001 |

Sodium per unit leaf area ~ ecotype * treatment + (1 | pair/accession)

| Effect | Estimate | SE | *z*-value | *P*-value |
| --- | --- | --- | --- | --- |
| Intercept(coastal water) | 0.293 | 0.5365 | 0.55 | 0.585 |
| Ecotype(inland) | -0.1365 | 0.7475 | -0.18 | 0.855 |
| Treatment(salt) | *4.*3394 | 0.5849 | 7.42 | <0.0001 |
| Treatment(salt):Ecotype(inland) | 2.6649 | 0.8355 | 3.19 | 0.0014 |

Succulence ~ ecotype * treatment + (1 | pair/accession)

| Effect | Estimate | SE | *z*-value | *P*-value |
| --- | --- | --- | --- | --- |
| Intercept(coastal water) | 0.0336 | 0.0014 | 24.08 | <0.0001 |
| Ecotype(inland) | -0.0076 | 0.002 | -3.85 | 0.0001 |
| Treatment(salt) | -0.0046 | 0.0011 | -4.09 | <0.0001 |
| Treatment(salt):Ecotype(inland) | -0.0044 | 0.0016 | -2.79 | 0.0053 |
