## Appendix S6 for "Foliar salt spray exclusion and tissue tolerance underlie local adaptation to oceanic salt spray"

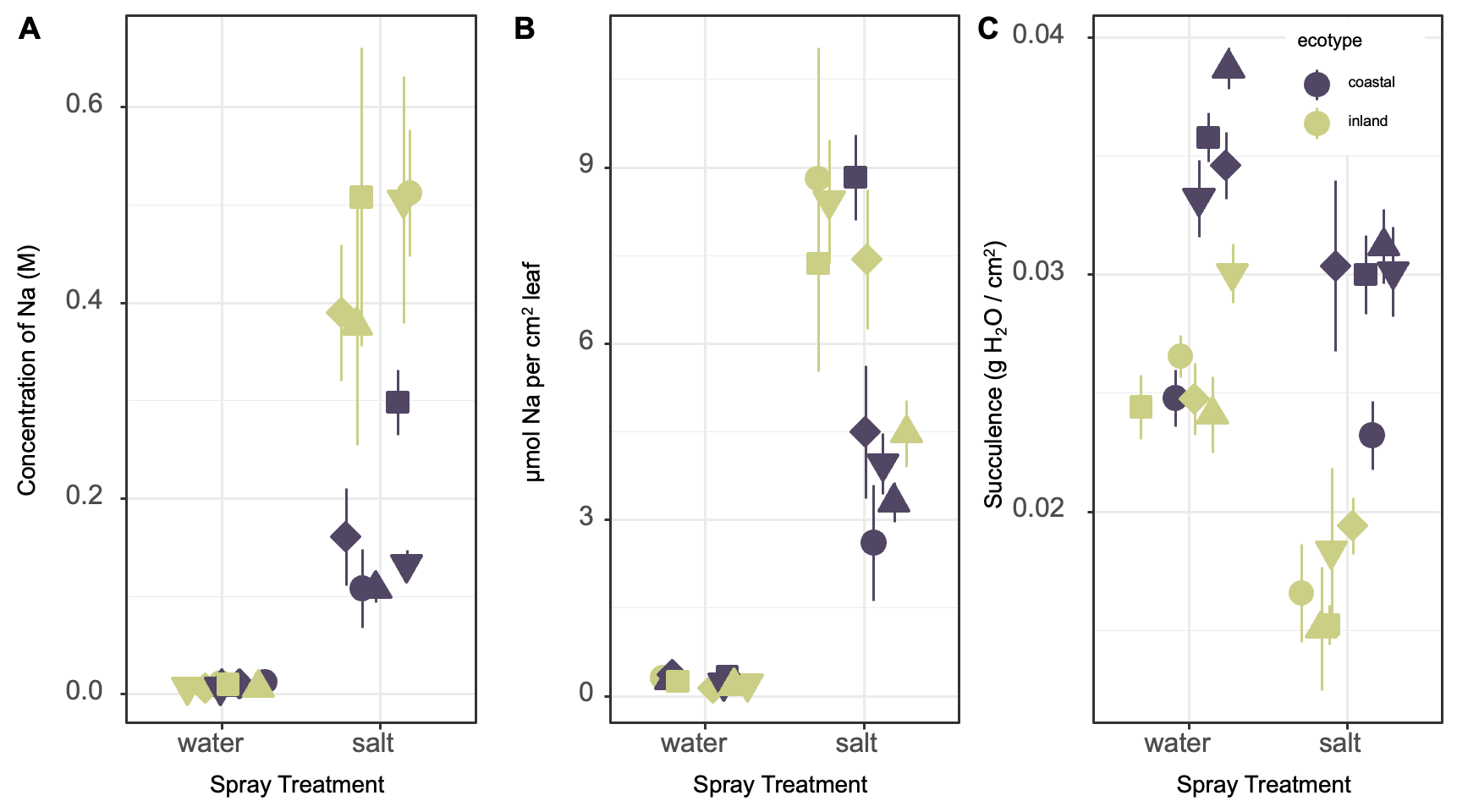

**Appendix S6.** Sodium and water levels in coastal and inland accessions, with means from four analyzed leaves for each accession under salt or water treatment. Error bars show standard errors of the mean. Shapes indicate accession, as shown in main text Figure 2. (A) Salt spray molarity for each ecotype and treatment, assuming all Na is in solution in the volume of water in the leaf at the time of sampling. (B) µmol of Na per unit leaf area for each ecotype and treatment. (C) Succulence as (fresh mass - dry mass) / leaf area, in grams H_2_O per cm^2^ leaf for each ecotype and treatment.
