## Appendix S7 for "Foliar salt spray exclusion and tissue tolerance underlie local adaptation to oceanic salt spray"

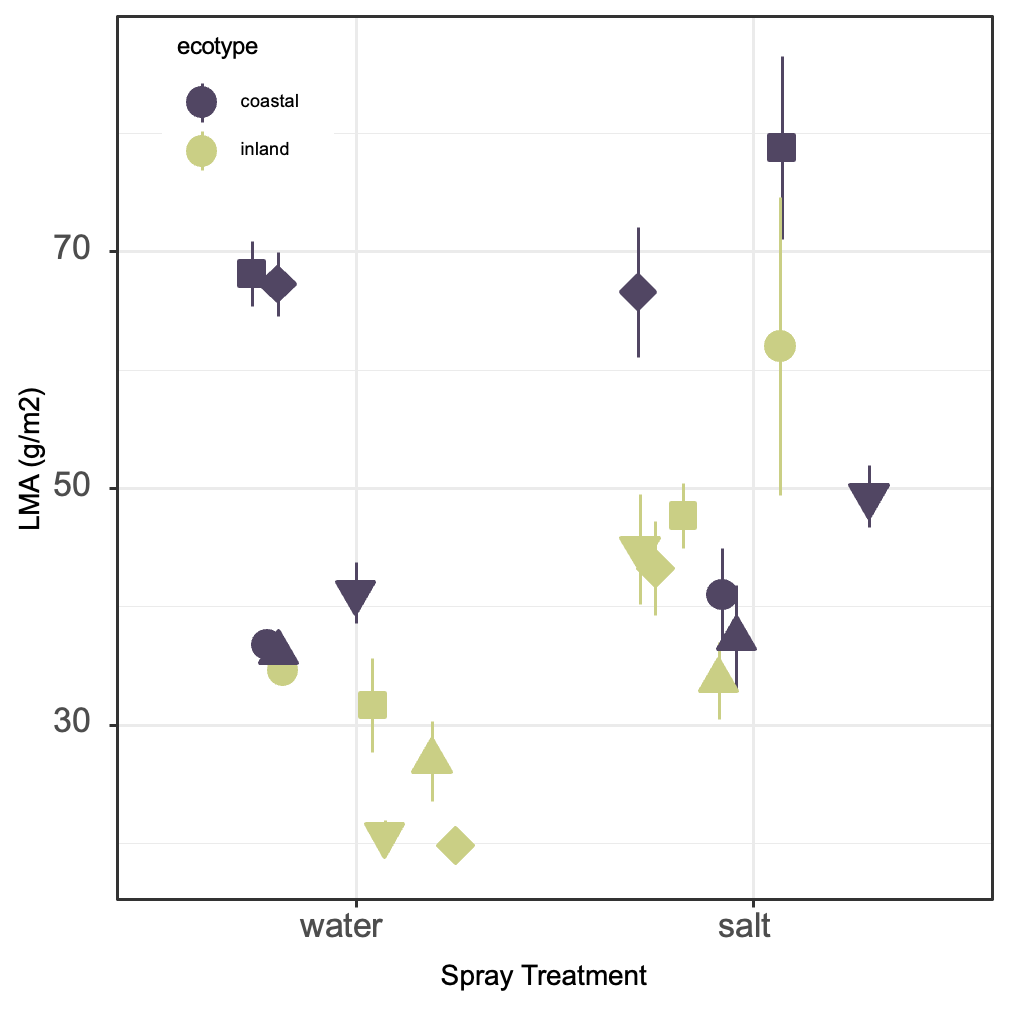

**Appendix S7.** Leaf mass per area (y-axis) in coastal and inland accessions after salt spray or control treatment, with means from four analyzed leaves for each accession under salt or water treatment (x-axis). Error bars show standard errors of the mean. Shapes indicate accession, as shown in main text Figure 2.
