## Appendix S8 for "Foliar salt spray exclusion and tissue tolerance underlie local adaptation to oceanic salt spray"

**Appendix S8.** Analysis of LMA (g m^-2^) under salt treatment with a general linear mixed model. Effect sizes are shown in the units of the response variable.

LMA ~ ecotype * treatment + (1 | pair/accession)

| Effect | Estimate | SE | *z*-value | *P*-value |
| --- | --- | --- | --- | --- |
| Intercept(coastal water) | 49.90 | 5.4432 | 9.17 | <0.0001 |
| Ecotype(inland) | -23.49 | 7.7125 | -3.05 | 0.0023 |
| Treatment(salt) | 4.70 | 2.798 | 1.68 | 0.093 |
| Treatment(salt):Ecotype(inland) | 14.81 | 3.998 | 3.70 | 0.0002 |
