## Appendix S9 for "Foliar salt spray exclusion and tissue tolerance underlie local adaptation to oceanic salt spray"

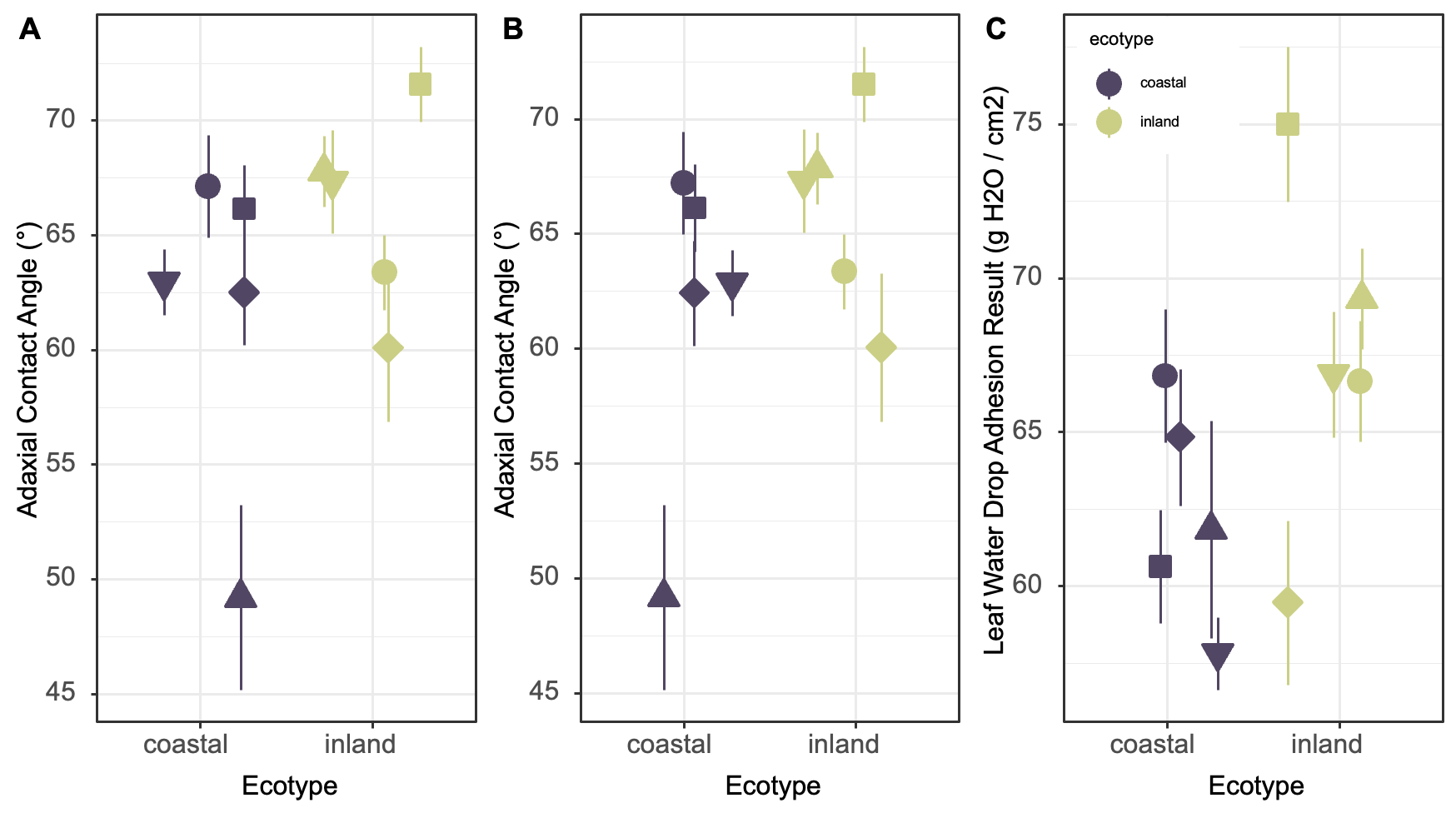

**Appendix S9.** Wettability traits (y-axis) in coastal and inland accessions (x-axis), with means from 15 analyzed leaves for each accession. Error bars show standard errors of the mean. Shapes indicate accession, as shown in main text Figure 2. (A) Contact angle in degrees of a drop of water placed on the adaxial surface, (B) Contact angle in degrees of a drop of water placed on the abaxial surface, (C) Result of leaf water drop adhesion assay in g H_2_O / cm^2^.
