## Appendix S10 for "Foliar salt spray exclusion and tissue tolerance underlie local adaptation to oceanic salt spray"

**Appendix S10.** Analyses of leaf wettability in coastal and inland accessions. General linear mixed models of contact angle of a drop of water on the adaxial and abaxial leaf surfaces (degrees), and leaf water drop adhesion assay (g H_2_O per cm^2^ of leaf area that adhered to a submerged leaf) as a function of ecotype and accession nested within ecotype. Wettable leaves have small contact angles and large leaf water drop adhesion assay results. Effect sizes are shown in the units of the response variable.

Adaxial contact angle ~ ecotype + (1 | pair/accession)

| Effect | Estimate | SE | *z*-value | *P*-value |
| --- | --- | --- | --- | --- |
| Intercept(coastal) | 61.56 | 2.40 | 25.66 | <0.0001 |
| Ecotype(inland) | 4.48 | 3.39 | 1.32 | 0.19 |

Abaxial contact angle ~ ecotype + (1 | pair/accession)

| Effect | Estimate | SE | *z*-value | *P*-value |
| --- | --- | --- | --- | --- |
| Intercept(coastal) | 62.42 | 1.89 | 32.96 | <0.0001 |
| Ecotype(inland) | 5.02 | 2.67 | 1.88 | 0.060 |

Leaf water drop adhesion assay result ~ ecotype + (1 | pair/accession)

| Effect | Estimate | SE | *z*-value | *P*-value |
| --- | --- | --- | --- | --- |
| Intercept(coastal) | 0.0133 | 0.0013 | 10.25 | <0.0001 |
| Ecotype(inland) | -0.0061 | 0.0017 | -3.61 | 0.0003 |
