## Appendix S11 for "Foliar salt spray exclusion and tissue tolerance underlie local adaptation to oceanic salt spray"

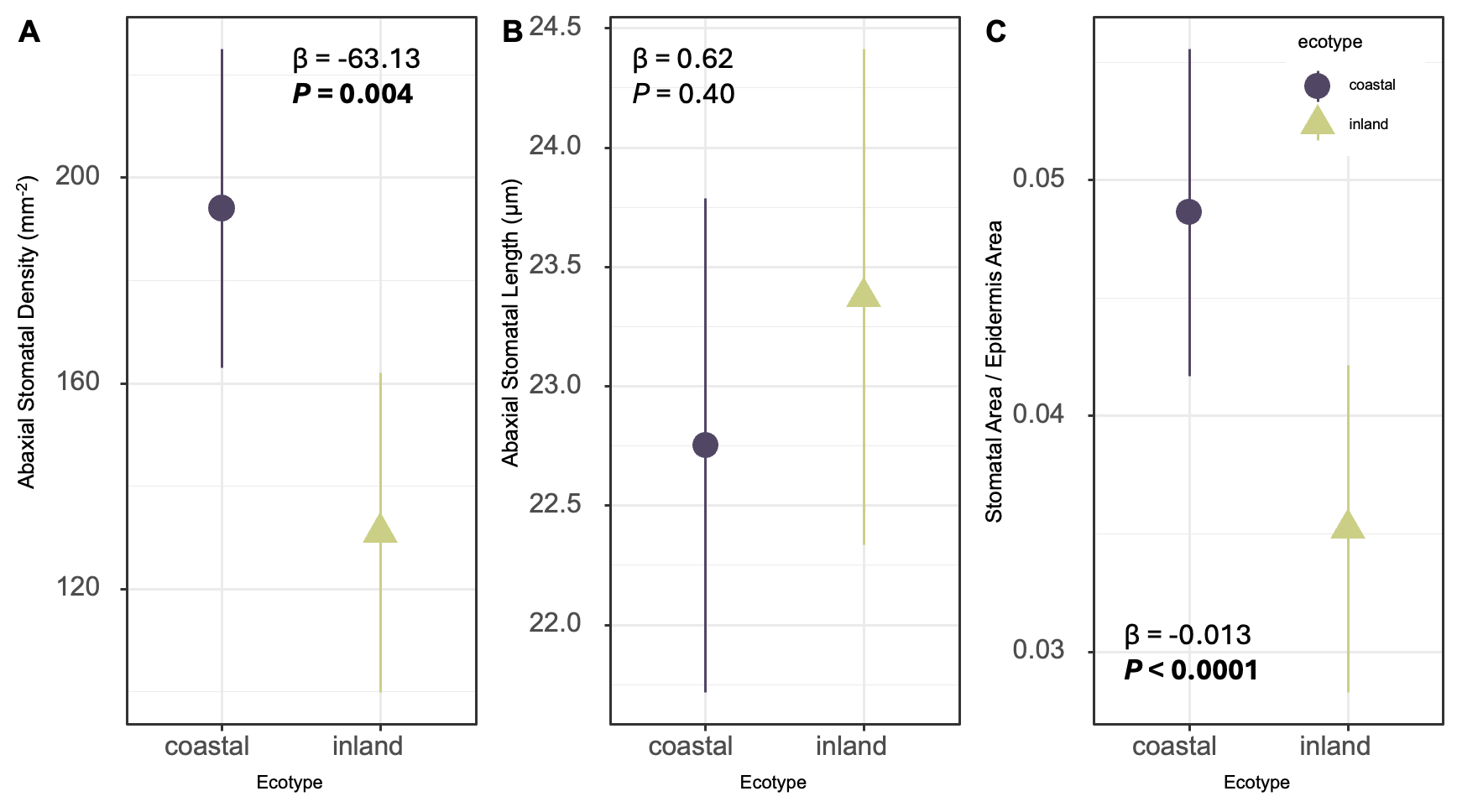

**Appendix S11.** Comparison of abaxial stomatal traits between ecotypes. Points indicate LSMs for each accession, with 15 individuals measured per accession, and error bars show confidence intervals. β and p-values indicate the effect and significance of inland ecotype on the trait from general linear mixed models. (A) Stomatal density in stomata per mm^2^ (y-axis) against ecotype (x-axis). (B) Stomatal length in µm (y-axis) against ecotype (x-axis). (C) Estimated fraction of epidermis area allocated to stomata (y-axis) against ecotype (x-axis).
