## Appendix S12 for "Foliar salt spray exclusion and tissue tolerance underlie local adaptation to oceanic salt spray"

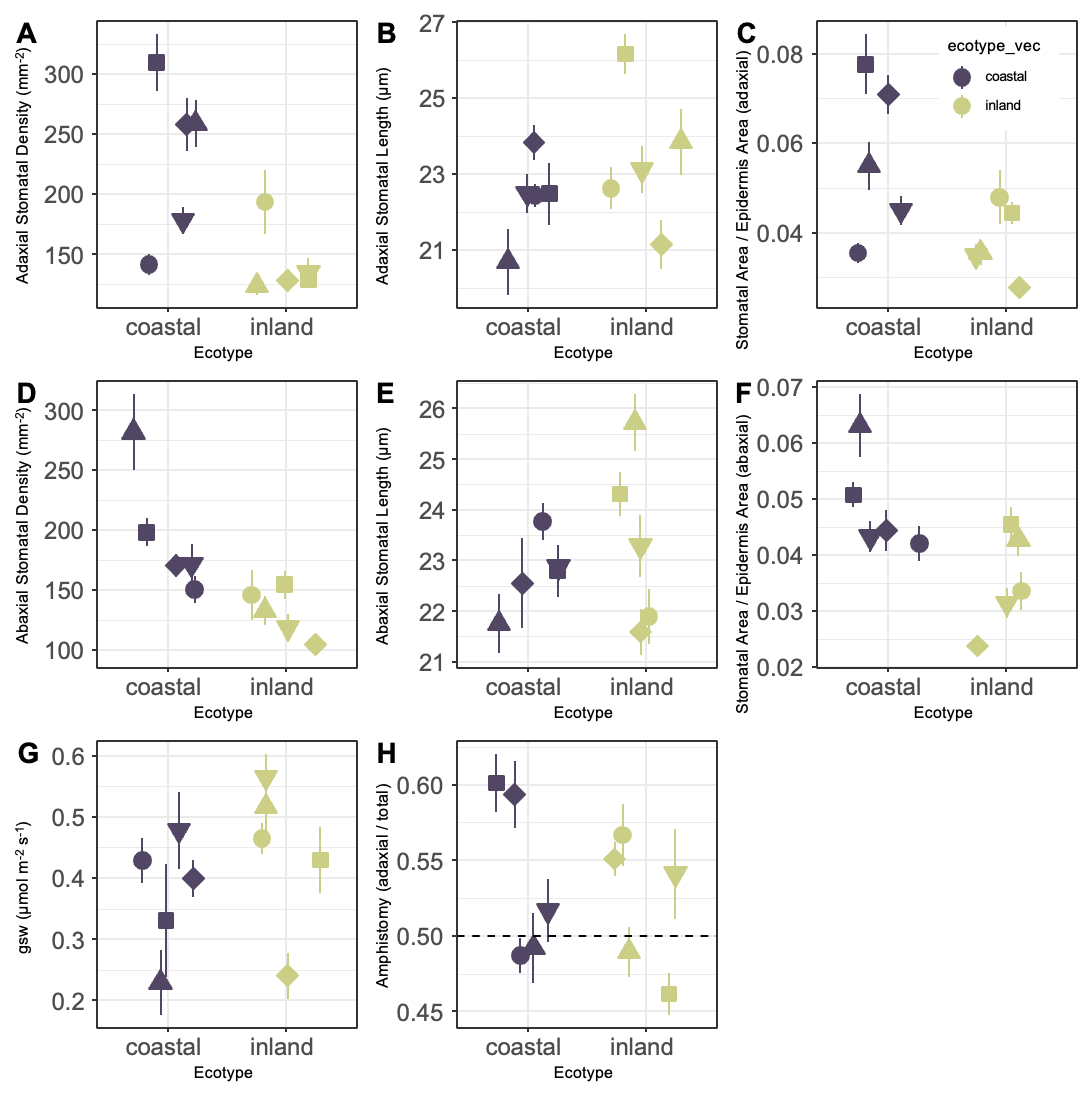

**Appendix S12.** Stomatal traits (y-axis) compared between coastal and inland accessions (x-axis). Points indicate means for each accession and error bars show standard errors of the mean. Traits were measured on 15 individuals from each accession for all traits except g_sw_, where 6 leaves sampled from 2-3 individuals were measured. (A) Adaxial stomatal density in stomata/mm^2^, (B) Adaxial stomatal length in μm, (C) Estimated fraction of adaxial epidermal area allocated to stomata, (D) Abaxial stomatal density in stomata/mm^2^, (E) Abaxial stomatal length in μm, (F) Estimated fraction of abaxial epidermal area allocated to stomata, (G) Stomatal conductance of adaxial surface as μmol m^-2^ s^-2^), amphistomy as adaxial stomatal count/(abaxial stomatal count + adaxial stomatal count), where dashed line at amphistomy = 0.5 corresponds to equal numbers of stomata on both leaf sides.
