## Appendix S13 for "Foliar salt spray exclusion and tissue tolerance underlie local adaptation to oceanic salt spray"

**Appendix S13.** Analyses of stomatal traits in coastal and inland accessions. General linear mixed models of stomatal density (number of stomata / mm^2^), stomatal length (μm), fraction of epidermis area allocated to stomata, stomatal conductance (as g_sw_, in mol m^-2^ s^-1^), and amphistomy (adaxial / total stomatal density). All traits except amphistomy are for the adaxial surface only.

Adaxial stomatal density ~ ecotype + (1 | pair/accession)

| Effect | Estimate | SE | *z*-value | *P*-value |
| --- | --- | --- | --- | --- |
| Intercept(coastal) | 229.00 | 20.89 | 10.96 | <0.0001 |
| Ecotype(inland) | -87.82 | 29.57 | -2.97 | 0.003 |

Adaxial stomatal length ~ ecotype + (1 | pair/accession)

| Effect | Estimate | SE | *z*-value | *P*-value |
| --- | --- | --- | --- | --- |
| Intercept(coastal) | 22.38 | 0.61 | 36.41 | <0.0001 |
| Ecotype(inland) | 0.996 | 0.87 | 1.14 | 0.25 |

Fraction adaxial epidermis allocated to stomata ~ ecotype + (1 | pair/accession)

| Effect | Estimate | SE | *z*-value | *P*-value |
| --- | --- | --- | --- | --- |
| Intercept(coastal) | 0.057 | 0.006 | 10.38 | <0.0001 |
| Ecotype(inland) | -0.019 | 0.008 | -2.43 | 0.015 |

Adaxial stomatal conductance ~ ecotype + (1 | pair/accession)

| Effect | Estimate | SE | *z*-value | *P*-value |
| --- | --- | --- | --- | --- |
| Intercept(coastal) | 0.374 | 0.045 | 8.24 | <0.0001 |
| Ecotype(inland) | 0.069 | 0.064 | 1.07 | 0.28 |

Amphistomy ~ ecotype + (1 | pair/accession)

| Effect | Estimate | SE | *z*-value | *P*-value |
| --- | --- | --- | --- | --- |
| Intercept(coastal) | 0.5382 | 0.0201 | 0.02 | <0.0001 |
| Ecotype(inland) | -0.0172 | 0.0285 | 0.028 | 0.55 |
