## Appendix S14 for "Foliar salt spray exclusion and tissue tolerance underlie local adaptation to oceanic salt spray"

**Appendix S14.** Analyses of abaxial stomatal traits in coastal and inland accessions. General linear mixed models of stomatal density (number of stomata / mm^2^), stomatal length (µm), and fraction of epidermis area allocated to stomata. Effect sizes are shown in the units of the response variable.

Abaxial stomatal density ~ ecotype + (1 | accession)

| Effect | Estimate | SE | z-value | *P*-value |
| --- | --- | --- | --- | --- |
| Intercept(coastal) | 194.04 | 15.6681 | 15.668 | <0.0001 |
| Ecotype(inland) | -63.13 | 22.1862 | 22.186 | 0.004 |

Abaxial stomatal length ~ ecotype + (1 | pair/accession)

| Effect | Estimate | SE | z-value | *P*-value |
| --- | --- | --- | --- | --- |
| Intercept(coastal) | 22.75 | 0.5237 | 0.524 | <0.0001 |
| Ecotype(inland) | 0.62 | 0.7418 | 0.742 | 0.40 |

Fraction abaxial epidermis allocated to stomata ~ ecotype + (1 | pair/accession)

| Effect | Estimate | SE | z-value | *P*-value |
| --- | --- | --- | --- | --- |
| Intercept(coastal) | 0.049 | 0.0035 | 0.004 | <0.0001 |
| Ecotype(inland) | -0.013 | 0.0028 | 0.003 | <0.0001 |
