## Appendix S15 for "Foliar salt spray exclusion and tissue tolerance underlie local adaptation to oceanic salt spray"

**Appendix S15.** Analysis of succulence in coastal and inland accessions, with 15 replicates per accession with no spray treatment. General linear mixed model of succulence (as (leaf fresh mass - leaf dry mass)/leaf area, in g H_2_O / cm^2^). Effect sizes are shown in the units of the response variable.

Succulence ~ ecotype + (1 | pair/accession)

| Effect | Estimate | SE | z-value | *P*-value |
| --- | --- | --- | --- | --- |
| Intercept(coastal) | 0.0289 | 0.0018 | 0.002 | <0.0001 |
| Ecotype(inland) | -0.0065 | 0.0026 | 0.003 | 0.012 |
