## Appendix S16 for "Foliar salt spray exclusion and tissue tolerance underlie local adaptation to oceanic salt spray"

**Appendix S16.** Analysis of tissue tolerance assay. General linear mixed models of days to the start of leaf disk necrosis and days to full leaf disk necrosis on salt and control media. Effect sizes are shown in the units of the response variable.

Days to start of necrosis ~ ecotype * treatment + (1 | pair/accession)

| Effect | Estimate | SE | *z*-value | *P*-value |
| --- | --- | --- | --- | --- |
| Intercept(coastal water) | 7.83 | 0.3243 | 24.16 | <0.0001 |
| Ecotype(inland) | 0.77 | 0.4626 | 1.66 | 0.097 |
| Treatment(salt) | -1.80 | 0.2851 | -6.30 | <0.0001 |
| Treatment(salt):Ecotype(inland) | -1.57 | 0.407 | -3.85 | 0.0001 |

Days to full necrosis ~ ecotype * treatment + (1 | pair/accession)

| Effect | Estimate | SE | *z*-value | *P*-value |
| --- | --- | --- | --- | --- |
| Intercept(coastal water) | 11.67 | 0.63 | 18.52 | <0.0001 |
| Ecotype(inland) | 0.39 | 0.90 | 0.44 | 0.66 |
| Treatment(salt) | -2.34 | 0.42 | -5.52 | <0.0001 |
| Treatment(salt):Ecotype(inland) | -0.59 | 0.61 | -0.98 | 0.33 |
